# Human brain microRNAs exhibit cell-type specificity and age dependence

**DOI:** 10.64898/2026.01.04.697535

**Authors:** Serafima Dubnov, Lihen Laski, Ido Zchut, Ofek Avidan, Estelle R. Bennett, David S. Greenberg, Adi Turjeman, Mor Nitzan, Iddo Paldor, Hermona Soreq

## Abstract

MicroRNAs play key roles in regulating brain processes ranging from neurogenesis to neurological disease and contribute to many cell-type-specific pathways. However, technical limitations of single-cell small RNA sequencing have confined most studies to bulk analyses of whole-tissue specimens, limiting our capacity to resolve the cell-type specificity of brain microRNAs and to study their roles in depth. To generate a comprehensive, cell-type-resolved atlas of microRNAs in the human brain, we isolated neurons, astrocytes, microglia and oligodendrocytes from fresh neurosurgery-derived brain samples and profiled their small RNA repertoires by RNA sequencing. The results revealed pronounced cell-type-associated differences in microRNA profiles, identified novel cell-type-specific microRNA markers and shed new light on the contribution of their originating genomic loci to cell-type specificity. We further characterized the cell-type dependence of microRNA strand preference and isomiR processing, and identified cell-type-dependent microRNA programs associated with brain ageing. Combined with an accompanying statistical tool for cell-type signal enrichment, our atlas provides a publicly available resource for cell-type-resolved microRNA analysis in the human brain.

## Introduction

MicroRNAs (miRs) are short non-coding RNA molecules, typically up to 22 nucleotides long, that can suppress gene expression through complementary binding to target mRNA transcripts^1,2^. miRs are well-established regulators of key brain processes^3,4^, including neurodevelopment^5–10^, synaptic plasticity^11–13^, apoptosis^14,15^, neuroinflammation^16^, neurodegeneration^17,18^ and epilepsy^4,19^. However, despite extensive evidence for their involvement in brain function, the cell-type-specific expression of brain miRs remains poorly characterized, limiting the functional interpretation of their regulatory roles in a cell-type-specific manner.

While single-cell RNA sequencing (scRNA-seq) has revolutionized transcriptomics by enabling the study of gene regulation at the resolution level of the individual cell, small non-coding RNAs (sncRNAs) are rarely captured by scRNA-seq that measures only a fraction of the transcriptome, leaving many lowly expressed transcripts, including miRs, undetected^20^. Additionally, most scRNA-seq protocols rely on poly(T) priming to capture polyadenylated mRNAs, whereas mature miRs are not polyadenylated^21^. Thus, sncRNA studies still tend to rely on bulk tissue samples that contain mixed combinations of cell types. Although several specialized protocols have been developed to measure miR expression at single-cell resolution, these approaches require dedicated methodology and are limited in the number of cells that can be profiled^22–26^. To our knowledge, no single-cell or cell-type-resolved miR dataset of the adult human brain currently includes neurons, astrocytes, microglia and oligodendrocytes in parallel. That being said, miRs do display cell-type-specific expression across multiple tissues^27,28^, including the brain^24^.

To address this knowledge gap, we constructed the first cell-type-resolved atlas of miR expression in the adult human brain, spanning neurons, microglia, astrocytes and oligodendrocytes. The atlas reveals both unique miR cell-type markers and shared, co-expressed miRs. Importantly, comparison with murine datasets^29–31^ shows substantial concordance in cell-type specificity, supporting evolutionary conservation. We find that many cell-type-enriched miRs are associated with quantitative trait loci (miR-QTLs) in cell-type-specific enhancers; that the specificity of intronic miRs largely tracks their host genes; and that microglial miR genes reside within microglia-specific regulatory regions. Based on our resource, we introduce a statistical tool that quantifies the degree of cell-type-specific marker enrichment in any given set of miRs. Applying this tool, we show that the bulk miR signatures underlying neurodegeneration, multiple sclerosis and ageing are enriched for neuronal, oligodendrocyte and microglial miRs, respectively. Finally, leveraging the wide donor age range, we identify age-associated miR profiles: neuron-enriched miRs targeting nervous system development genes decrease with age, in contrast to neuron- and astrocyte-enriched miRs targeting hallmark ageing pathways that increase, implicating miRs in cell-type-specific processes of brain ageing.

Our study provides a unique resource for investigating cell-type-specific expression patterns of miRs in the adult human brain. Both the dataset and the accompanying cell-type enrichment tool are available for interactive usage on our website (https://brain-visualization.vercel.app/).

## Results

### Characterizing the cell-type specificity of human brain miRs

To establish a cell-type-resolved resource of brain microRNAs, we collected 39 brain tissue samples (17 females, 22 males; ages 37–85; Supplementary Table 1) obtained during neurosurgical procedures, by collecting the brain parenchyma along the surgical access route to the tumor. Only patients with non-infiltrative pathologies were included in this study, so that the collected specimens can be considered to represent healthy brain tissue rather than tumor tissue (Methods). We then disassociated the collected specimens to single nuclei with a surrounding perinuclear cytoplasmic fraction using the NuNeX protocol^32^ and separated cell types by fluorescence-activated cell sorting (FACS; Figure 1a). Nuclei were stained with DAPI for the identification of single cells rather than debris or doublets, and the cell type-characteristic markers NeuN, GFAP and IBA1 were used to identify neurons, astrocytes and microglia, respectively (Supplementary. Fig. 1a-h; Methods).

**Figure 1.**
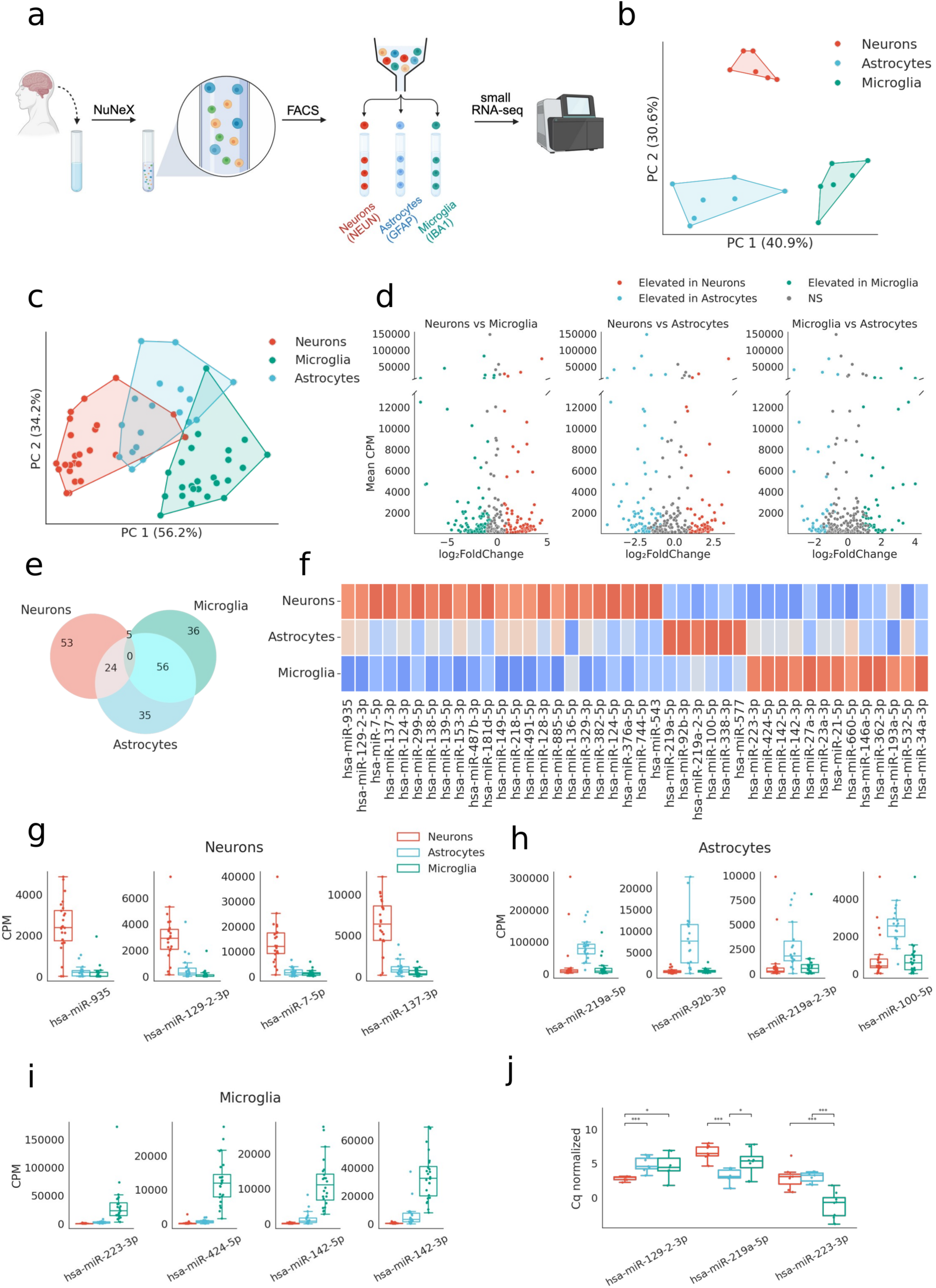
Cell-type-specificity of human brain miRs. a) Graphical representation of the pipeline used to produce the cell-type-resolved resource of small non-coding RNAs from live human brain. Created with BioRender. b) PCA of GAPDH-normalized Cq values of cell-type mRNA markers (neuronal: SNAP25, SYN1 SYNPR; astrocytic: AQP4, ATP13A4, GJA1, SLC1A2, SLCO1C; microglial: ALOX5AP, CD68, CD74) across all samples, colored by cell type. Convex hulls outline each cell-type cluster. c) Scatter plot of Principal Component Analysis (PCA) applied to miR profiles of cell-type-specific populations, obtained with small RNA-seq. Convex hulls outline each cell-type cluster. d) Volcano plots showing dream pairwise DE analysis, comparing each pair of cell types. e) Venn diagram showing the overlaps between the elevated miRs in each cell type. f) Heatmap of mean cell type z-scores of marker miRs for each cell type, defined as significantly DE miRs with a log-fold change greater than 2, relative to both other cell types (Methods). g) Box plot showing expression levels of the top five neuron-enriched miRs (Methods). h) Box plot as in f for astrocyte-enriched miRs. i) Box plot as in f for microglia-enriched miRs. j) Box plots of qPCR Cq values, normalized to hsa-let-7a-5p, for three cell-type-enriched miRs, one per cell type, measured across all three cell types. *p ≤ 0.05, **p ≤ 0.01, ***p ≤ 0.001 (non-parametric Mann–Whitney U test).

Total RNA was extracted from the sorted populations, and a subset of samples was used to validate their cell type identities by qPCR measurement of additional cell type markers. NeuN-positive nuclei expressed the neuronal markers SNAP25, SYN1 and SYNPR (Supplementary Fig. 1k), GFAP-positive nuclei expressed the astrocytic markers AQP4, ATP13A4, GJA1, SLC1A2 and SLCO1C1 (Supplementary Fig. 1l), and IBA1-positive nuclei expressed the microglial markers ALOX5AP, CD68 and CD74 (Supplementary Fig. 1m). Principal component analysis (PCA) of Cq values normalized to the GAPDH gene yielded distinct separation of the cell-type groups, confirming that the sorted populations correspond to the intended cell types with minimal overlap (Figure 1b).

To compare miR levels across brain cell types, we profiled 24 NeuN-positive, 24 IBA1-positive, and 18 GFAP-positive samples with small RNA-sequencing (Supplementary Table 1, Methods). To account for statistical dependencies in the data, such as repeated measurements from the same patient and global effects of batch, sex, age and pathology type, we used the dream framework^33^. Variance partitioning revealed cell type as the strongest predictor of miR levels (mean explained variance 18.3%), followed by patient ID (mean explained variance 12%; Supplementary Fig. 2a). By contrast, batch, pathology type, sex and age explained only minor fractions of variance (mean explained variances 3.0%, 0.7%, 0.5%, and 1.7%, respectively; Supplementary Fig. 2a), highlighting the strong cell-type-specific regulation of miRs and validating that the tumor pathology had minimal influence on the miR profiles.

Importantly, the miR profiles clustered according to their cell type label: k-means clustering resulted in sample division that closely matched the annotated cell types (Figure 1c, Supplementary Fig. 2b), yielding a normalized mutual information (NMI) score of 0.8 (Methods). In contrast, the NMI score between the k-means clusters and batch labels was 0.1, implying that the miR profiles clustered by cell type rather than by batch (Supplementary Fig. 2c). To identify cell-type-enriched miRs we performed DE analysis in three pairwise comparisons (neurons vs. microglia, neurons vs. astrocytes and microglia vs. astrocytes), correcting for batch, pathology type, sex, age and patient ID (Figure 1d; Supplementary Tables 2-4). Out of 280 miRs expressed above a defined threshold (Methods), 209 were significantly DE in one of the pairwise cell type comparisons (Figure 1d). Specifically, we identified 191 DE miRs between neurons and microglia, 163 between neurons and astrocytes, 96 between microglia and astrocytes (Figure 1d).

Through pairwise comparisons across the three cell types, we identified miRs significantly elevated in more than one cell type (Figure 1e): 24 shared by neurons and astrocytes, 56 by astrocytes and microglia, and 5 by neurons and microglia (Supplementary Table 5). After excluding these overlaps we identified 53, 36, and 35 miRs uniquely enriched in neurons, microglia and astrocytes, respectively (Figure 1e).

The top miR markers for each cell type were defined as those with a log-fold change greater than 2 compared to both other cell types. We identified 23, 13 and 6 top miR markers in neurons, microglia and astrocytes (Figure 1f, Table 1, Supplementary Table 6, Methods). For neurons, the top 4 (by fold change) miRs were hsa-miR-935, hsa-129-2-3p, hsa-miR-7-5p and hsa-miR-137-3p (Figure 1g). Their neuronal identities are well supported in the literature: miR-7 regulates neuronal death and glutamatergic signaling^34,35^, miR-137 is critical for neural development^36^, and miR-935 and miR-129-2-3p are both elevated in epileptic brains and plasma^37^. Among astrocytic miRs, hsa-miR-92b-3p was enriched, and has previously been linked to astrocytes in murine brain^38^ and shown to regulate glioblastoma development in the human brain^39^ (Figure 1h). Top microglial miRs included the hsa-miR-142 family, previously associated with microglia in murine brain^38^ and linked to neuroinflammation in murine models^40,41^ (Figure 1i). We further validated several of the identified miR markers by qPCR, confirming enrichment of hsa-miR-129-2-3p in neurons, hsa-miR-219a-5p in astrocytes and hsa-miR-223-3p in microglia (Figure 1j). miR Cq values were normalized to hsa-let-7a-5p, which showed uniform expression across cell types in our atlas (Supplementary Fig. 2d). All unique cell-type markers identified in this study are listed in Table 1.

**Table 1.** Cell-type-specific miR markers in the adult human brain.

| miRNA | Verified and conserved | Host gene (cell-type zscore) | Ageing |
| --- | --- | --- | --- |
| *Neurons* |  |  |  |
| miR-935 |  |  |  |
| miR-129-2-3p |  |  |  |
| miR-7-5p |  |  |  |
| miR-137-3p |  |  |  |
| miR-124-3p |  |  |  |
| miR-299-5p | Yes<sup>30,31</sup> |  |  |
| miR-138-5p | Yes<sup>30,31</sup> |  |  |
| miR-139-5p | Yes<sup>30</sup> | PDE2A (1.63) |  |
| miR-153-3p |  |  |  |
| miR-487b-3p | Yes<sup>31</sup> |  |  |
| miR-181d-5p |  |  |  |
| miR-149-5p |  | GPC1 (1.06) |  |
| miR-218-5p | Yes<sup>31,32</sup> |  |  |
| miR-491-5p | Yes<sup>30,31</sup> |  |  |
| miR-128-3p |  |  |  |
| miR-885-5p |  | ATP2B2 (1.24) |  |
| miR-136-5p | Yes<sup>31</sup> |  | Increased |
| miR-329-3p | Yes<sup>31,32</sup> |  | Decreased |
| miR-382-5p | Yes<sup>30,31</sup> |  |  |
| miR-124-5p |  |  | Increased |
| miR-376a-5p | Yes<sup>30,31</sup> |  |  |
| miR-744-5p | Yes<sup>30</sup> | MAP2K4 (1.50) |  |
| miR-543 | Yes<sup>30,31</sup> |  |  |
| *Microglia* |  |  |  |
| miR-223-3p | Yes<sup>31</sup> |  | Increased |
| miR-424-5p |  |  |  |
| miR-142-5p | Yes<sup>31</sup> |  |  |
| miR-142-3p | Yes<sup>31</sup> |  |  |
| miR-27a-3p |  |  |  |
| miR-23a-3p |  |  |  |
| miR-21-5p | Yes<sup>31</sup> |  |  |
| miR-660-5p |  | CLCN5 (1.91) |  |
| miR-146a-5p | Yes<sup>31</sup> |  |  |
| miR-362-3p | Yes<sup>31</sup> | CLCN5 (1.91) |  |
| miR-193a-5p |  |  |  |
| miR-532-5p | Yes<sup>31</sup> | CLCN5 (1.91) | Increased |
| miR-34a-3p | Yes<sup>31</sup> |  |  |
| *Astrocytes* |  |  |  |
| miR-92b-3p |  |  |  |
| *Astrocytes & Oligodendrocytes* |  |  |  |
| miR-219a-5p | Yes<sup>31</sup> (Ast) |  |  |
| miR-219a-2-3p | Yes<sup>31,32</sup> (Ast,Glia) |  |  |
| miR-100-5p | Yes<sup>31</sup> (Ast) |  |  |
| miR-338-3p | Yes<sup>31</sup> (Ast) | AATK (1.95, Olig) |  |
| miR-577 |  | UGT8 (2.00, Olig) |  |
| *Oligodendrocytes* |  |  |  |
| miR-33b-5p |  | SREBF1 (1.52) |  |
| miR-181b-5p |  |  |  |
| miR-338-5p |  | AATK (1.95) |  |
| miR-574-3p |  | FAM114A1 (1.27) |  |
| miR-151b |  |  |  |
| miR-151a-5p |  | PTK2 (1.53) |  |
| miR-625-5p |  | FUT8 (1.80) |  |
| miR-151a-3p |  | PTK2 (1.53) |  |
| miR-181a-5p |  |  |  |
| miR-330-3p |  | EML2 (1.37) |  |
| miR-625-3p |  | FUT8 (1.80) |  |

miRs that are enriched in two cell types but depleted in the third are of particular interest as well, possibly pointing to shared cell-type specific functions (Supplementary Table 5). We found elevated levels of hsa-miR-9, a regulator of synaptic plasticity and memory^42^, in both neurons and astrocytes (Supplementary Fig. 2e). The hsa-miR-221/222 family, involved in neuroinflammation^43–45^, was enriched both in neurons and microglia (Supplementary Fig. 2f). Finally, glial miRs enriched in both microglia and astrocytes relative to neurons included hsa-miR-32-5p, which has been linked to neuroinflammation mediated by both glial cell types^46,47^ (Supplementary Fig. 2g).

Markers are grouped by the cell type in which they are enriched (for neurons, microglia and astrocytes: padj < 0.05 and log2FC ≥ 2 in all pairwise comparisons; for oligodendrocytes: padj < 0.05 and log2FC ≥ 1.5 in the comparison of oligodendrocytes vs. the mean of all other cell types). All miRs are human (hsa-). The “Verified and conserved” column indicates whether the marker was verified as enriched in the same cell type in an external dataset (referenced), unless a different cell type is specified in parentheses (Ast, astrocytes; Glia, all glial types). The “Host gene” column lists the host genes of intronic miRs together with the mean z-score of the corresponding cell type from a single-cell atlas (Lee et al., 2024). In the “Ageing” column ‘Increased’ indicates increased abundance with age (bioIB metagene 3 score > 0.01) and ‘Decreased’ indicates decreased abundance with age (bioIB metagene 1 score > 0.01).

### FACS isolation of oligodendrocytes derived from live human brain tissue

To enhance the depth of our cell-type-resolved resource, we also profiled microRNAs from the fourth major brain cell type – oligodendrocytes. We used the well-established oligodendrocyte-specific OLIG2 antibody, which requires an additional permeabilization step to allow antibody access to the nucleus. This necessitated separate staining and FACS sorting steps (Figure 2a, Methods). As in the original protocol (Figure 1a), single cells were identified by DAPI (Supplementary Fig. 3a), and OLIG2-positive nuclei were gated based on thresholds defined from non-stained controls (Supplementary Fig. 3b,c, Methods). Altogether, we profiled miRs from 14 sorted OLIG2-positive populations (5 females and 9 males, Supplementary Table 1), using small RNA sequencing.

**Figure 2.**
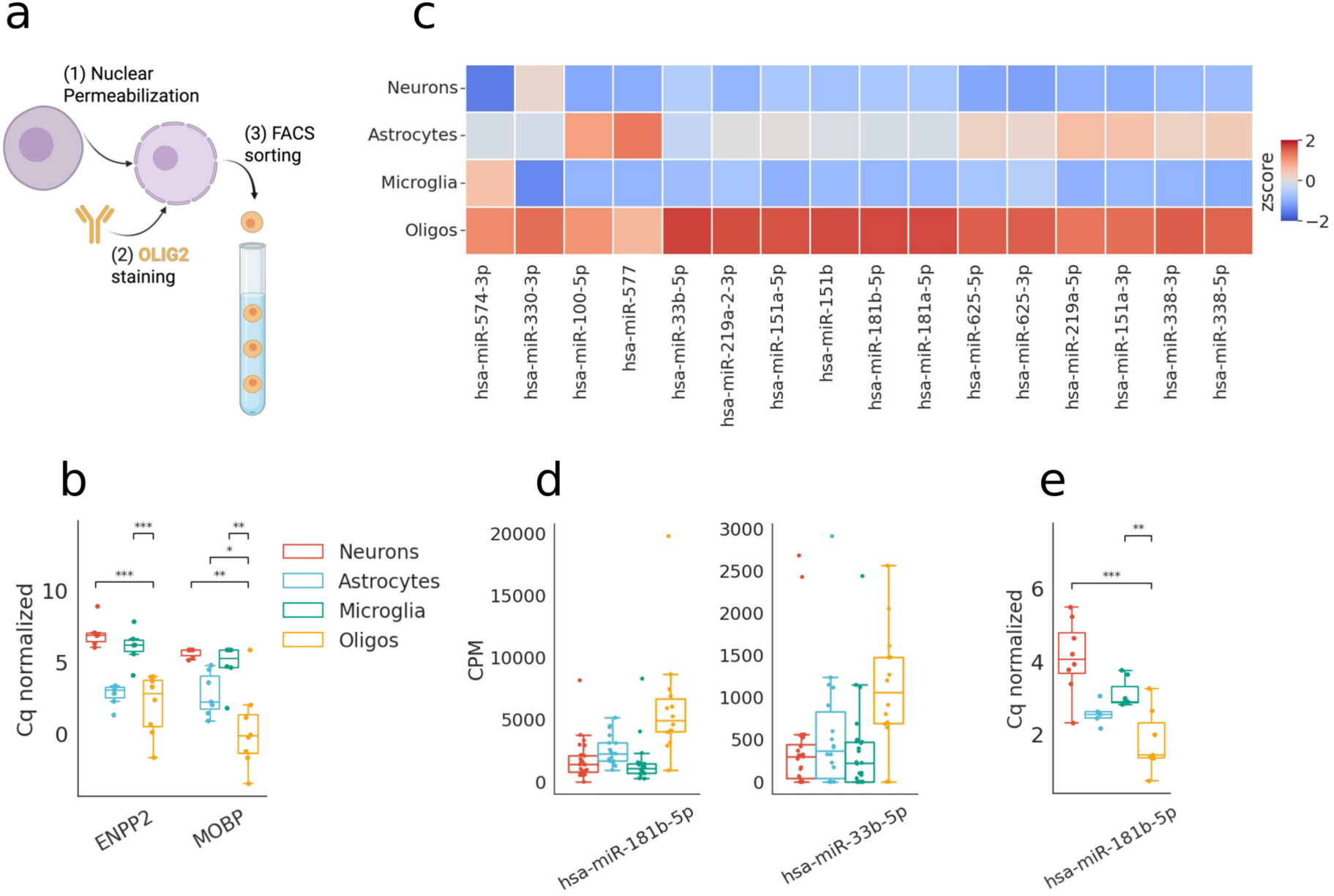
Identifying oligodendrocyte-enriched miRs. a) Graphical representation of the FACS sorting pipeline used to isolate oligodendrocytes from live human brain specimens. Created with BioRender. b) Box plot showing qPCR Cq values for ENPP2 and MOBP normalized to GAPDH, measured in all four cell types. c) Heatmap showing z-scores of 16 miRs whose levels were significantly elevated in oligodendrocytes (log2 fold change > 1.5, adjusted p ≤ 0.05), compared to all other cell types. d) Boxplots showing the levels of two oligodendrocyte marker miRs (log2 fold change > 2; adjusted p ≤ 0.05; Methods). CPM - counts per million. e) Box showing qPCR Cq values for hsa-miR-181b-5p normalized to hsa-let-7a-5p, measured in all four cell types. For b) and e) *p ≤ 0.05, **p ≤ 0.01, ***p ≤ 0.001 (non-parametric Mann–Whitney U test).

As with the other cell types, we used qPCR to measure enrichment of the oligodendrocyte biomarkers ENPP2 and MOBP in the OLIG2-positive fraction compared with neurons, microglia, and astrocytes (Figure 2b). However, the OLIG2-positive population also featured increased levels of the neuronal marker SNAP25, the astrocyte markers AQP4 and ATP13A4 and the microglial marker ALOX5AP (Supplementary Fig. 3d-g). Therefore, the OLIG2-positive group represents a mixed, albeit oligodendrocyte-enriched population.

The mixed identity of the OLIG2-positive population justified a different statistical approach to identify the oligodendrocyte-enriched miRs. To find such miRs, we applied the dream framework^33^ by comparing the OLIG2-positive population to the averaged profile of the three other cell types while correcting for batch, pathology type, sex, age and participant ID (Methods). This analysis revealed 16 miRs that were significantly enriched in oligodendrocytes relative to all other cell types (Figure 2c, Table 1, Supplementary Table 6). Notably, these included hsa-miR-219a and hsa-miR-338, both known to promote myelin repair^48^, and hsa-miR-23, which enhances oligodendrocyte differentiation and myelin synthesis^49^. We also identified hsa-miR-181a-5p, previously implicated in oligodendrocyte maturation^50^ and suggested as a potential epigenetic biomarker of multiple sclerosis^51^. To identify the oligodendrocyte miR markers we further excluded miRs with log-fold change lower than 2, as well as those that appeared as markers of other cell types. This resulted in two miRs that were uniquely enriched in the oligodendrocyte population: hsa-miR-181b-5p and hsa-miR-33b-5p (Figure 2d). The miR-33 family is associated with lipid metabolism in the brain^52^, and the miR-181b family is implicated in myelin-assisted axon regeneration^53^. The oligodendrocyte enrichment of hsa-miR-181b-5p was also validated by qPCR (Figure 2e). Having extracted the oligodendrocyte miR signature alongside neuron-, astrocyte-, and microglia-specific profiles, we could thereby extend our resource to encompass all four major brain cell types.

### miR cell-type specificity shows evolutionary conservation

The existing data on miR cell-type specificity in humans is limited. Datasets derived from primary cell cultures do not faithfully recapitulate their in vivo counterparts^54^, while studies of single purified cell types cannot compare relative enrichment between cell types and consequently cannot establish miR cell-type specificity^55^. The few existing cell-type-resolved miR datasets are restricted to the developing human brain^24,38^, and therefore do not capture the miR repertoires of mature glial and neuronal populations in the mature brain. In contrast, our atlas is unique in its ability to simultaneously compare miR signatures across four distinct cell types of the adult human brain. To validate our findings and to assess the evolutionarily conservation of the cell-type specificity we identified, we compared our findings with three murine datasets of varying cell-type resolution and coverage (Figure 3a).

**Figure 3.**
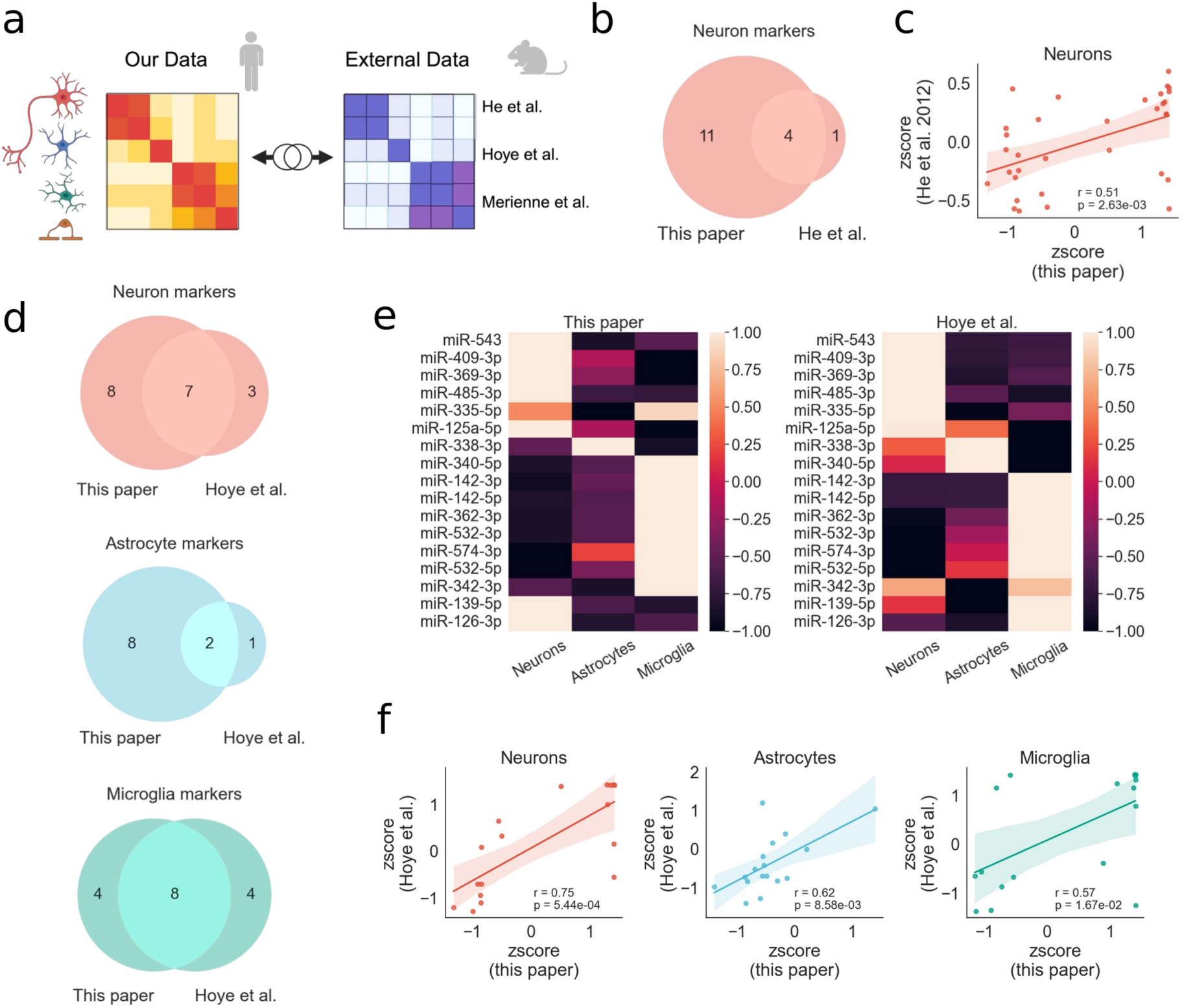
Evolutionary conservation of miR cell-type specificity. a) Schematic overview of the analysis performed in this section. Cell-type specificity of the miR profiles generated in this study was compared with three independent datasets from murine brain, each profiling miRs in isolated cell types. Created with BioRender. b) Venn diagram comparing miRs enriched in neurons in this study with miRs significantly elevated in neuronal cell types relative to bulk whole-cortex and cerebellum samples in He et al.^29^ (Mann–Whitney test, logFC > 1, FDR < 0.05). Only conserved miRs detected in both studies are shown. c) Scatterplot showing normalized expression levels (z-scores) of shared neuronal miRs in our dataset and in He et al.^29^ d) Venn diagrams comparing the numbers of cell-type-specific miRs in neurons, astrocytes, and microglia identified in our dataset with miRs significantly elevated in the corresponding cell types in Hoye et al.^30^, based on the original analysis reported in that study. Only conserved miRs detected in both studies are shown. e) Heatmaps showing mean cell-type-wise normalized expression levels (z-scores) of 17 shared miRs in our dataset and in Hoye et al.^30^ f) Scatterplots showing the correlation of the z-scores shown in (e), separately for each cell type. Pearson’s *R* and corresponding *p* values are shown on the plots.

First, we compared our findings with those of He et al.^29^, who introduced miR tagging and affinity purification (miRAP) and applied it to compare miR signatures across neuronal subtypes of the cortex and cerebellum against the bulk signatures of the corresponding whole regions. Thirty-three conserved miRs were detected in both datasets, four of which were significantly upregulated in neurons in both studies (Figure 3b). Notably, the neuronal miR markers identified in our study were significantly enriched in the neuronal populations of He et al., relative to non-neuronal miRs (Supplementary Fig. 4a). Conversely, the microglial miR markers identified in our study were predominantly depleted in those neuronal populations relative to non-microglial miRs (Supplementary Fig. 4b). Comparing normalized expression of the overlapping miRs across neuronal samples in the two datasets further revealed a significant correlation (Pearson R = 0.51; p = 0.003; Figure 3c).

Second, we compared our results with those of Hoye et al.^30^, who applied the same miRAP approach to profile miR signatures across three cell types: neurons, astrocytes and microglia. We found 7, 2 and 8 overlapping markers of neurons, astrocytes and microglia, representing 39%, 18% and 50% out of all conserved miR markers that were detected in both studies (Figure 3d). More generally, the relative expression of the overlapping miRs was highly correlated within each cell type (Figure 3e, f), supporting the validity of our data and demonstrating conserved cell-type associations among the shared miRs.

Finally, we compared our data with that of Merienne et al.^31^, who isolated neurons, astrocytes and microglia from mouse brain by laser capture microdissection (LCM) using transgenic mice expressing GFP in all three populations. Of the 8 neuronal miR markers identified in their study and detected in our data, 5 were also enriched in neurons in our atlas (Supplementary Fig. 4c). Moreover, miR-219a-2-3p and miR-34a-5p, reported as glia-enriched in their study, were upregulated in our atlas in both astrocytes and microglia (Supplementary Fig. 4d).

Together, these findings indicate that miR cell-type specificity is evolutionarily conserved, thus further validating the cell-type-specific miR profiles reported here.

### The genomic locus of a miR predicts its cell-type specificity

We next asked whether the cell-type specificity of miRs is determined by their genomic location. Using the FANTOM5 integrated atlas of miRs and their promoters^56^, we found that many of the miR cell-type markers identified in our study are encoded within the introns of protein coding genes (Figure 4a). We therefore compared the cell-type specificity of these miRs with that of their host genes. To assess the cell-type-specific expression of the host genes we used single-cell RNA-sequencing data from postmortem dorsolateral prefrontal cortex (DLPFC) of neurotypical donors^57^ (Methods). 70 of the intronic-miR host genes were represented in this dataset (Supplementary Table 7), and they exhibited strong cell-type specificity: their expression profiles separated cells by type with an NMI of 0.8 (Figure 4b). We then performed a correlation analysis between the relative cell-type expression of intronic miRs in our atlas and that of their host genes in the scRNA-seq data. A significant positive correlation was observed in all cell types except astrocytes, which showed a positive but non-significant trend, indicating that the cell-type specificity of intronic miRs is linked to that of their host genes (Figure 4c). The host genes of the intronic miR cell-type markers identified in this study are listed in Table 1, together with the z-scores of the corresponding cell types from the single-cell RNA-seq data^58^.

**Figure 4.**
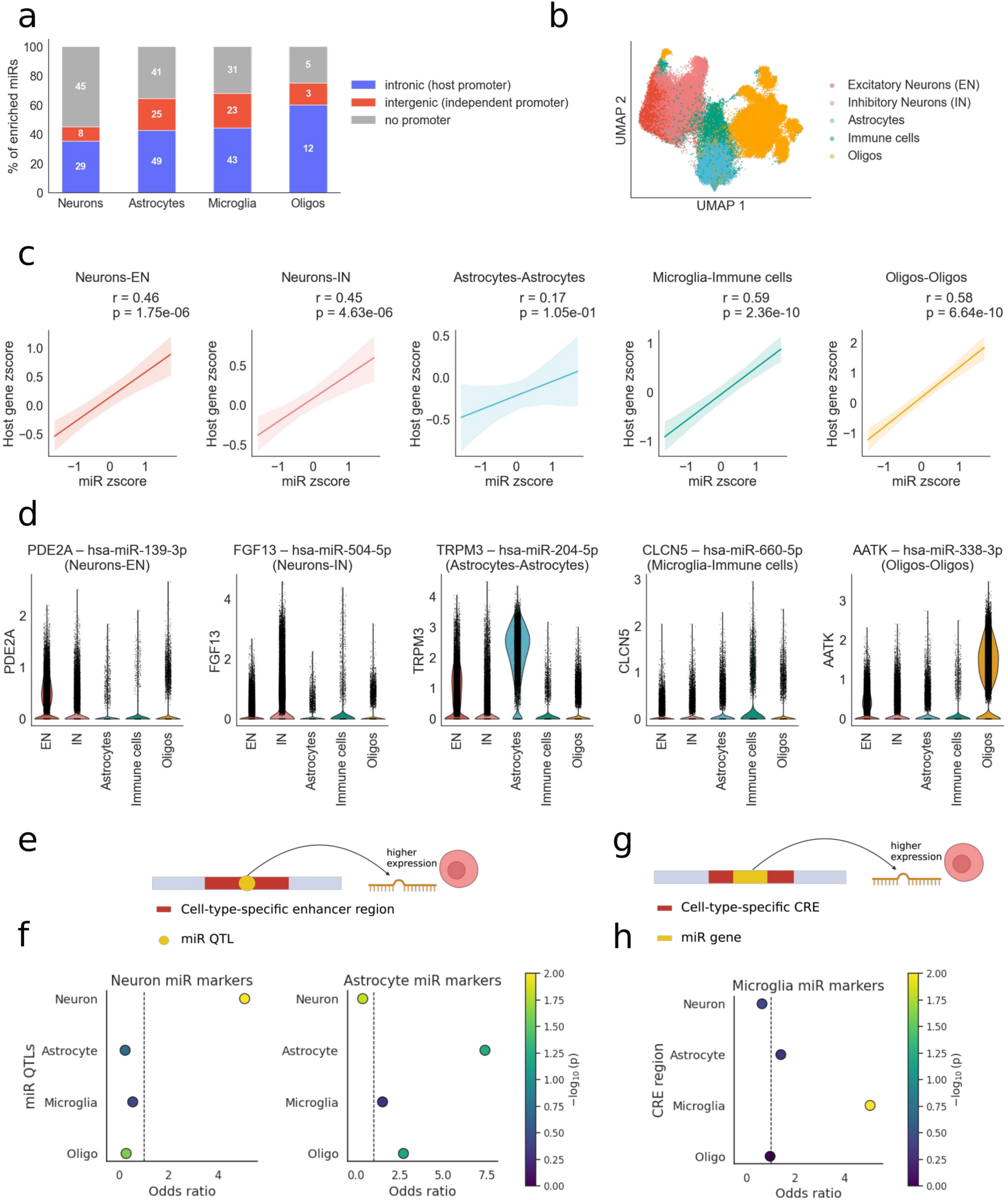
The genomic locus of a miR predicts its cell-type specificity. a) Stacked bar plot showing the proportions of intronic miRs (regulated by their host gene promoter), intergenic miRs (regulated by an independent promoter), and miRs lacking an annotated promoter among the cell-type-enriched miRs identified here. b) Uniform Manifold Approximation and Projection (UMAP) of the scRNA-seq dataset from postmortem DLPFC of neurotypical controls^57^, based on the expression of the 70 intronic-miR host genes. c) Correlation plots between the mean relative expression (z-scores) of intronic miRs and their host genes, per cell-type pair (our atlas vs. scRNA-seq): neurons vs. excitatory neurons; neurons vs. inhibitory neurons; microglia vs. immune cells; oligodendrocytes vs. oligodendrocytes; astrocytes vs. astrocytes. Pearson’s *R* and corresponding *p* values are shown above each plot. d) Violin plots of scRNA-seq expression (per cell type) for the host genes of the miR–host pairs with the highest mean z-score. e) Schematic representation of a miR-QTL (yellow) residing within a cell-type-specific enhancer region (red), thereby contributing to the cell-type-specific expression of associated miRs. Created with BioRender. f) Dot plots showing Fisher’s exact test results (alternative = two-sided) for the association between miRs enriched in neurons (left) and in astrocytes (right), and the associated miR-QTLs residing within cell-type-specific enhancer regions. Y-axis depicts cell type, X-axis positions reflect the odds ratio of each association, and dot colors depict -log10 p-values (unadjusted). g) Schematic representation of a miR gene (yellow) located within a cell-type-specific cis regulatory element (CRE) (red) that contributes to cell-type-specific expression of the miR. Created with BioRender. h) Dot plot showing Fisher’s exact test results (alternative = two-sided) for the association between miRs enriched in microglia and the cell-type-specific cis-regulatory elements (CREs). Y-axis depicts cell type, X-axis positions reflect the odds ratio of each association, and dot colors depict -log10 p-values (unadjusted).

We also identified the host–miR pairs that maximized the mean z-score in the corresponding cell types (Figure 4d). For example, PDE2A, enriched in excitatory neurons (log2FC = 2.3, padj ≈ 0), is the host of hsa-miR-139-3p, enriched in neurons in our atlas. FGF13, enriched in inhibitory neurons (log2FC = 3.3, padj ≈ 0), hosts hsa-miR-504-5p, also enriched in neurons. CLCN5, elevated in immune cells (log2FC = 2.4, padj = 6 × 10⁻¹¹⁹), hosts hsa-miR-660-5p, enriched in microglia. AATK, enriched in oligodendrocytes (log2FC = 2.9, padj ≈ 0), hosts hsa-miR-338-3p, also enriched in oligodendrocytes. Finally, TRPM3, enriched in astrocytes (log2FC = 3.7, padj ≈ 0), hosts hsa-miR-204-5p, enriched in astrocytes in our atlas.

miR expression is also subject to cis-regulation by genetic variation at miR quantitative trait loci (miR-QTLs)^59^. We therefore hypothesized that cell-type-enriched miRs may also be regulated by QTLs located within cell-type-specific promoter or enhancer regions (Figure 4e; Supplementary Table 8). Leveraging the previously annotated miR-QTLs^59^, we identified 31 neuron-, 20 microglia-, 21 astrocyte-, and 4 oligodendrocyte-enriched miRs regulated by 152, 113, 108, and 39 QTLs, respectively. To examine whether these QTLs overlap with cell-type-specific regulatory regions we mapped their genomic coordinates to cell-type-specific promoter and enhancer annotations^60^. We then applied Fisher’s exact test^61^ to statistically evaluate the association between these cell-type-enriched miRs and their QTLs residing in cell-type-specific genomic regions. Neuronal miRs indeed showed significantly enriched QTLs within neuron-specific enhancers (p = 2 × 10⁻⁵; Figure 5f, left panel) and a similar, though weaker, enrichment trend was observed between astrocytic miRs and miR-QTLs residing in astrocyte-specific enhancer regions (p = 0.0575; Figure 4f, right panel). The association between microglial miRs and miR-QTLs in microglia-specific enhancers, as well as between oligodendrocyte-enriched miRs and miR-QTLs in oligodendrocyte-specific enhancers, did not reach statistical significance (Supplementary Fig. 5b, Supplementary Table 8). Similarly, no significant associations were observed between miR levels and cell-type-specific promoter regions (Supplementary Table 8). These results collectively suggest that miR-QTLs residing in cell-type-specific enhancer regions may contribute to the genetic regulation of cell-type-specific miR expression.

**Figure 5.**
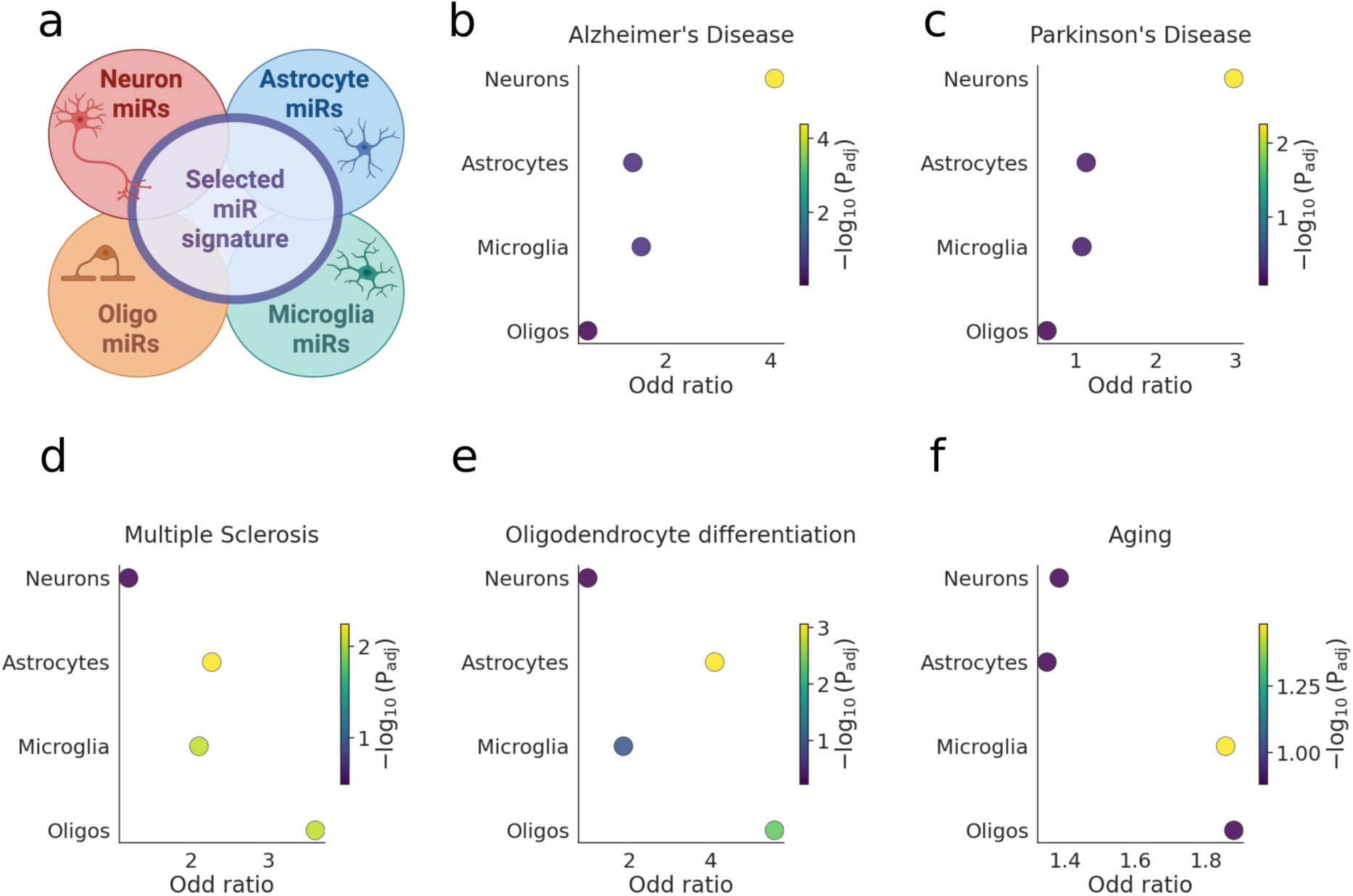
Cell type enrichment in miR lists of interest. a) Graphical representation of the pipeline used to deconvolve miR lists of interest to specific cell type signals. Cell-type enrichment for a given list is calculated by applying the Fisher’s exact test to the overlaps between the cell type miR markers identified in this study, and the input miR list. Thus, every analysis consists of four independent statistical tests (for the association with each of the four cell types), corrected for multiple comparisons. Created with BioRender. b-f) Fisher’s exact test (*alternative = greater*) results for deconvolution of (b) AD^66^, (c) PD^67^, (d) MS^68^, (e) oligodendrocyte differentiation^70^ and (f) brain ageing^71^ miR signatures. Reported *p*-values were adjusted for multiple testing using false discovery rate control.

We next investigated whether the genomic loci of cell-type-enriched miRs overlap with cell-type-specific cis-regulatory (CRE) regions^62^ (Figure 4g). Using a single-cell chromatin accessibility atlas of the adult human brain^62^ we mapped the genomic coordinates of cell-type-enriched miRs to chromatin regions differentially accessible in diverse cell types, and quantified their associations using Fisher’s exact test (Methods). Microglial miRs showed the strongest association within microglia-specific CRE regions (p = 0.0012; Figure 4h, Supplementary Fig. 5d, Supplementary Table 9), suggesting that chromatin accessibility contributes to the cell-type-specificity of microglial miRs.

### Cell-type-specific strand selection

Mature miRs are produced from either the 5’ or the 3’ arm of the miRNA precursor hairpin, and differences in their arm preference might have a direct impact on cellular function^63^. We therefore searched for miRNA precursors with cell-type-dependent arm preference.

First, we compared the levels of mature miRs produced from the 5’ and 3’ arms of the same precursor within each cell type. In every cell type, the levels of the two arms were significantly anti-correlated (Supplementary Fig. 6a). To quantify arm preference and compare it across cell types, for each precursor we calculated an arm preference index (API):

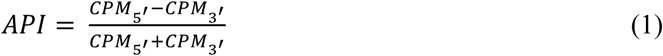

An API of +1 or −1 indicate exclusive 5’-arm or 3’-arm usage, respectively. We computed the API for each precursor in each cell type and compared API values across cell types per precursor (Supplementary Fig. 6b). The strength of arm preference (|*API*|) was negatively correlated with the variance of API across cell types (Supplementary Fig. 6c), indicating that the stronger a precursor’s bias toward one arm, the more likely that arm remains dominant across all cell types.

Nevertheless, we identified 5 miRNA precursors that switch their dominant arm in a cell-type-dependent manner, characterized by elevated API variance and sign switching across cell types (Supplementary Fig. 6d, Supplementary Table 10). Specifically, hsa-miR-590 predominantly produces the mature 3’-arm miR in astrocytes but the 5’-arm miR in neurons; hsa-miR-193b produces the 3’-arm miR in microglia and the 5’-arm in neurons; hsa-miR-10401 produces the 3’-arm miR in oligodendrocytes and microglia but the 5’-arm in astrocytes and neurons; hsa-miR-485 predominantly produces the 5’-arm in astrocytes but both arms in the other cell types; and hsa-miR-369 produces the 3’-arm miR in microglia but both arms in the other cell types (Supplementary Fig. 6d, Supplementary Table 10).

### Cell-type-dependent isomiR processing

IsomiRs – transcripts that differ from the canonical miR sequence by one or a few nucleotides – have been reported to exert distinct functions^64,65^. To probe 5′ and 3′ isomiR differences between brain cell types, we quantified the abundances of miR transcripts carrying 5′- and 3′-end truncations and overhangs (Methods). We first compared the normalized levels of each isomiR family with those of the corresponding canonical transcripts, computing Pearson correlations between z-scored isomiR and canonical miR expression within each cell type. IsomiR levels were generally highly correlated with those of the canonical miRs (Supplementary Figure 7a), with astrocytes showing the strongest coupling across isoform classes (mean R = 0.91), followed by microglia (mean R = 0.83), neurons (mean R = 0.76), and oligodendrocytes (mean R = 0.66). To identify individual miRs with differential isoform usage across cell types we next compared relative isomiR abundance, defined as the ratio of each isomiR to its canonical transcript levels, between cell types using ANOVA with Tukey post-hoc tests (Methods). This analysis yielded three significant 3′ truncation isomiRs (hsa-miR-3929, hsa-let-7a-5p, hsa-miR-23a-3p) and six 3′ overhang isomiRs (hsa-miR-222-3p, hsa-let-7c-5p, hsa-miR-32-5p, hsa-miR-190a-5p, hsa-let-7d-5p, hsa-let-7a-5p; Supplementary Figure 7b). Notably, hsa-let-7c-5p, previously implicated in stroke (Ni et al., 2015), showed elevated 3′ overhang isomiR abundance in neurons relative to glial cell types (Supplementary Table 11).

### Measuring cell-type enrichment in candidate miR sets

By revealing cell-type-specific miR markers, our atlas can serve to deconvolve cell-type-specific signals within a miR set of interest (Methods). Here, we provide a statistical framework that calculates the cell-type enrichment of a list of input miRs, such as a list of DE miRs from a bulk RNA-seq study. Specifically, we calculate the fraction of each cell type’s marker miRs present in the input list and assess the significance of that association using Fisher’s exact test with multiple comparison correction^61^ (Figure 5a). To maximize coverage, we include overlapping markers; thus, if an input miR is enriched in more than one cell type, it contributes to the enrichment of each type.

We applied this framework to a range of miR sets derived from small RNA-seq in studies of brain-related conditions and pathologies (Supplementary Table 12). The signature of 110 DE miRs identified in the AD study^66^ showed significant enrichment for neuronal markers (Figure 5b). Similarly, 125 miRs altered in PD brains^67^ were also enriched for neuronal markers (Figure 5c). This suggests that the miR signatures associated with neurodegeneration mainly reflect neuronal processes. In contrast, the 63 miRs associated with multiple sclerosis^68^ (MS) were predominantly enriched in oligodendrocyte-specific miRs, but also showed significant contributions of astrocyte and microglia markers, consistent with the role of glial-driven neuroinflammation in MS^69^ (Figure 5d). A curated set of 17 miRs functionally linked to oligodendrocyte differentiation was enriched for the oligodendrocyte and astrocyte signatures, further validating our framework^70^ (Figure 5e). We also examined a murine ageing-related miR study comprising 476 miRs^71^, and although this set contained markers from all major brain cell types, only microglial enrichment was statistically significant, consistent with the original study that highlighted microglial miR involvement in brain ageing^71^ (Figure 5f). In summary, our cell-type-resolved brain miR atlas enabled the assignment of miR signatures to specific brain cell types, providing a powerful tool for interpreting bulk miR profiling studies.

### Elucidating age-dependent miRs with bioIB

Although multiple miRs have been proposed as key regulators of ageing-related processes in the mouse brain^71–73^, age-associated miR expression patterns in the human brain remain poorly characterized. Our cohort spans a wide age range (37 – 85 years of age) and is composed of individuals with no cognitive impairment, thus providing a unique opportunity to study miR dynamics during healthy ageing of the brain. Previous studies have shown that miRs often change in coordinated clusters with correlated expression patterns^71,74^. We therefore hypothesized that analyzing age-associated miR clusters could reveal ageing-related signatures. To this end, we employed bioIB^75^, our recently developed framework for identifying maximally informative, signal-aware representations of high-dimensional data (Methods).

Samples were divided into two age groups: younger (<65 years) and older (≥65 years). To avoid bias arising from differential cell-type composition, we subsampled the data to include an equal number of samples from each of the four cell types in both age groups, selecting the six samples with the highest RNA integrity numbers (RINs) per cell type and age group. This resulted in a balanced dataset of 48 samples (six samples per cell type per age group). Using age group as the signal of interest, hierarchical bioIB partitioned miRs into three major metagenes: metagene 1, enriched in younger individuals; metagene 2, a neutral metagene; and metagene 3, enriched in older individuals (Figure 6a). To rule out the possible confounding effect of pathology type, we compared the normalized expression of these metagenes between the brain metastases and meningioma groups and found no significant differences (Supplementary Figure 8). Because metagene 2 did not show age-specific enrichment, subsequent analyses focused on the two age-associated metagenes (1 and 3).

**Figure 6.**
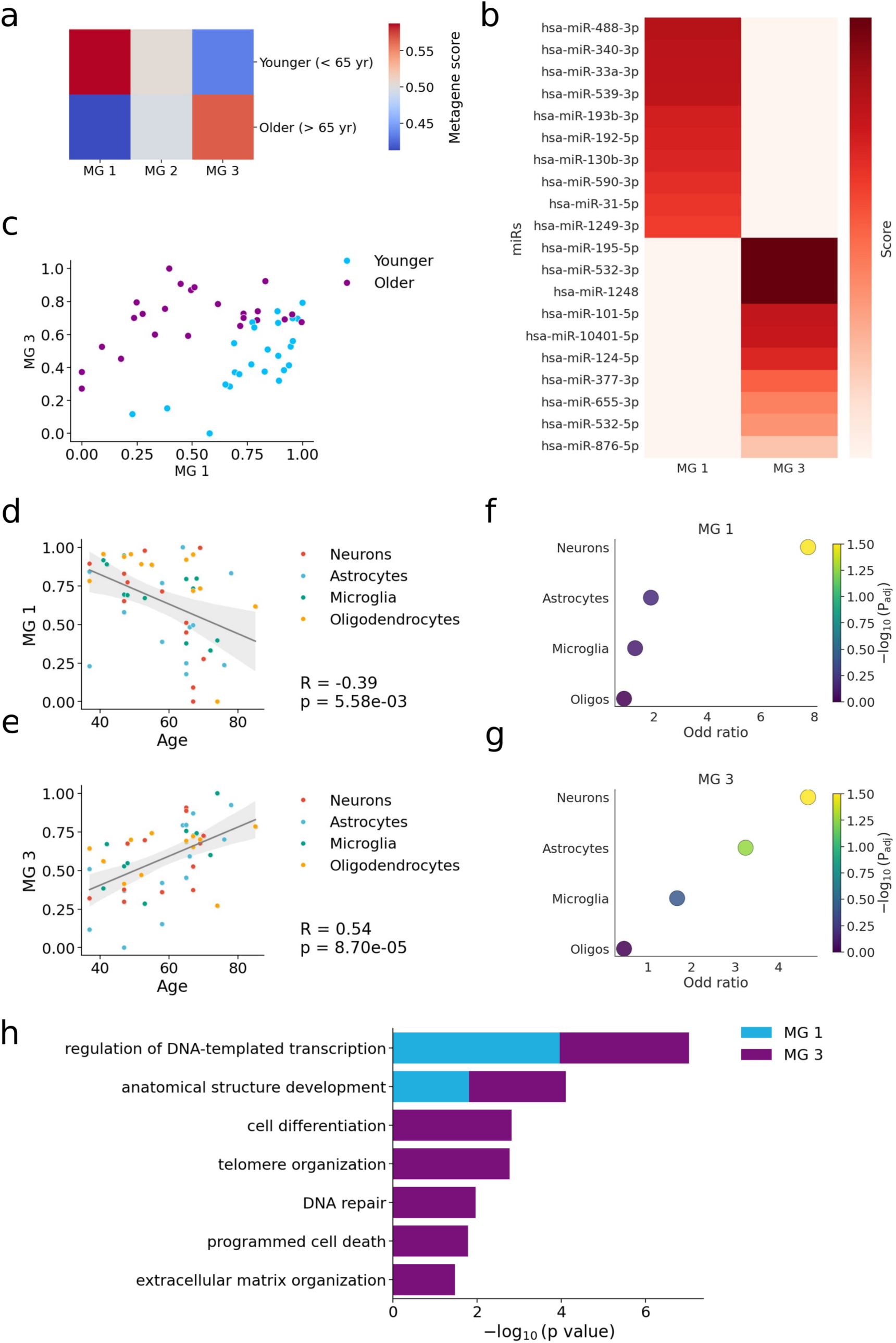
Determining age-dependent miRs with bioIB. a) Heatmap showing bioIB metagene scores across age groups. Scores are normalized within each metagene, with higher values indicating higher relative metagene expression in the corresponding age group. Metagene 1 is enriched in younger individuals (bioIB metagene score = 0.6 in the younger group); metagene 2 is a neutral metagene (score = 0.5 in both groups); metagene 3 is enriched in older individuals (score = 0.6 in the older group). b) Heatmap showing bioIB metagene scores for the top 10 representative miRs per metagene. Scores are normalized within each metagene, with higher values indicating a stronger contribution of a given miR to that metagene. c) Scatter plot of samples included in the bioIB analysis (younger, n = 24; older, n = 24), showing normalized expression of metagene 1 (x-axis) versus metagene 3 (y-axis). Points are colored by age group. d) Scatter plot showing normalized metagene 1 expression as a function of age, with points colored by the cell-type origin of each sample. Shaded region indicates the 95% confidence interval of the regression fit (bootstrap-estimated). e) Same as (d) for metagene 3. f) Dot plot showing cell-type enrichment in the list of representative miRs for metagene 0. g) Same as (f) for metagene 3. h) Bar plot showing the top GO Slim categories enriched among predicted targets of miRs representing each bioIB metagene. Enrichment p-values (X-axis) correspond to the lowest adjusted p-value among GO terms mapping to each GO Slim category.

BioIB enables direct mapping between each metagene and its representative miRs (Figure 6b; Supplementary Table 13). The top miR representing metagene 1 was hsa-miR-488-3p, recently implicated in regulating ageing hallmark genes^76^. Another prominent member of this metagene, hsa-miR-193b-3p, has been shown to attenuate neuroinflammation^77,78^. In contrast, the top miRs representing metagene 3, hsa-miR-195-5p and hsa-miR-532-3p, have both been reported to exert neuroprotective effects in the context of brain injuiry^79–81^.

Clustering samples based on the expression of metagenes 1 and 3 separated individuals primarily by age group (NMI = 0.35), rather than by cell type (NMI = 0.09), indicating that the identified metagenes capture age-related rather than cell-type-specific variation (Figure 6c; Methods). Consistent with this, metagene 1 exhibited a significant negative correlation with age (Spearman R = −0.39, adjusted p-value < 0.0001), contrasting metagene 3 that showed a significant positive correlation with age (Spearman R = 0.54, adjusted p-value < 0.0001) (Figure 6d, e).

For each metagene we next defined the most representative miRs as those featuring metagene representation scores higher than 0.01, yielding 17 miRs for metagene 1 and 19 miRs for metagene 3 (Supplementary Table 13). Metagene 1, decreasing with age, featured one neuronal marker, hsa-miR-329-3p, whereas metagene 3, increasing with age, featured two neuronal markers, hsa-miR-136-5p and hsa-miR-124-5p, and two microglial markers, hsa-miR-223-3p and hsa-miR-532-5p (Table 1, Supplementary Table 13). Furthermore, we assessed cell-type enrichment with our deconvolution framework (Figure 5a, Methods). Both metagenes appeared as significantly enriched for neuronal miRs, while metagene 3 also showed significant enrichment for astrocytic miRs (Figure 6f, g). Finally, for each metagene we extracted the predicted and experimentally validated targets of its representative miRs using miRTarBase^82–84^, and applied the Gene Ontology^85,86^ (GO) enrichment analysis to elucidate the common targeted pathways (Methods; Supplementary Table 14). GO revealed shared enrichment for transcriptional regulation and central nervous system development processes (Figure 7h, Supplementary Table 14). Metagene 3, positively correlated with age, also included miRs targeting ageing-related pathways such as telomere organization, DNA repair, and apoptosis^87^. Collectively, these results suggest that coordinated, cell-type-specific miR programs are associated with the healthy ageing of the human brain.

## Discussion

In this study we have established the first cell-type-resolved atlas of miR expression profiles in the adult human brain. The identification of both unique and shared miR markers across four major brain cell types revealed that brain miR profiles are highly cell-type-specific. Furthermore, we showed that miRNA cell-type specificity is strongly shaped by genomic context: intronic miRs largely inherit the cell-type specificity of their host genes, while other miRs are influenced by miR-QTLs and by their own location within cell-type-specific enhancers and other cis-regulatory elements. We also characterized strand preference across cell types: for most miRs, 5′- or 3′-arm dominance was stable across all cell types, though a subset showed cell-type-dependent arm preference. Likewise, isomiR levels were generally highly correlated with those of the canonical miRs, with a few miRs displaying slight cell-type dependency. Finally, using our atlas, we were also able to identify cell-type-enriched miR profiles that are associated with ageing. These included miRs with reported neuroprotective functions that target genes involved in hallmark ageing-related processes.

We have also introduced a statistical framework for cell-type enrichment analysis of miR lists of interest. Since no robust, commercial scRNA-seq technology currently enables the capture of small RNA profiles, most studies of brain small RNAs are based on bulk RNA-seq. In this context, our atlas offers an opportunity to deconvolve cell-type signals from bulk-derived miR profiles. Once scalable single-cell small RNA sequencing becomes feasible, our resource may also serve as a reference resource for cell type annotation, providing validated miR markers for neuronal and glial populations.

Our atlas was generated from fresh human brain tissue, which, on the one hand, represents its unique strength by avoiding postmortem transcriptional artifacts. On the other hand, although our specimens were obtained from the parenchyma surrounding the tumor, not the tumor itself, we cannot exclude the possibility that tumor-related processes influenced the microRNA profiles of the adjacent tissue. To minimize this effect we used specimens obtained only during resections of non-invasive brain tumors. All the resected tumors were non-infiltrative and external to the brain, originating from the meninges (in meningioma) or from non-neural tissue (in metastases). Our previous work on this cohort^88^ demonstrated that these samples are transcriptionally closer to postmortem human brain tissues than to tumor-associated profiles. Additionally, statistical analysis in this paper demonstrated minimal effect of tumor pathology on miR profiles. Both these points suggest minimal tumor effects on our results.

A further limitation of our study is the lower yield of small RNAs obtained from the NuNeX^32^ protocol compared to whole-cell extraction. Because NuNeX isolates nuclei together with a surrounding perinuclear cytoplasmic fraction, our profiling predominantly captures cytoplasmic small RNAs located in the perinuclear compartment. Nevertheless, miRs are known to localize to the nucleus, endoplasmic reticulum and mitochondria^89^, all of which are retained in the NuNeX method.

In summary, this study represents an effort to decipher the microRNA landscape of human brain cells. Future work could extend this atlas to finer cellular states such as neuronal subtypes^24^ or disease-associated glial populations^90–92^, and integrate emerging single-cell small-RNA sequencing technologies to achieve higher resolution. Both our cell-type-resolved miR atlas, and the miR cell-type deconvolution tool are publicly available for interactive exploration on our portal (https://brain-visualization.vercel.app/).

## Materials and methods

### Tissue samples

Specimens were obtained during neurosurgical resection of non-neural lesions such as meningiomas and brain metastases. They were taken along the surgical access route to the tumor, from the surrounding, uninfiltrated brain parenchyma. In selecting patients for our cohort we followed strict exclusion criteria and did not include patients with preoperative evidence of damaging brain processes such as infarcts or bleeding, or patients who were operated on for infiltrative pathologies such as low-grade gliomas, primary CNS lymphoma or glioblastoma. Thus, although neurosurgical procedures are by definition performed to treat brain pathology, and the brain surrounding the pathology itself could be affected via edema, perilesional microvascular changes, microbleeds or other microenvironment changes, the excised samples themselves were not cancerous.

After the surgical extraction, tissue samples (average size of 3 x 5 mm) were fixed in 4% formaldehyde (Sigma-Aldrich, HT501128) for 30 minutes, washed with PBS (Merck, D8537) and stored at 4°C in RNAlater® (Merck, R0901) for 1-3 days until homogenization.

### NuNeX^32^

Brain sections were homogenized in 5 mL nuclear isolation buffer (10mM Tris pH 8.0, 25 mM KCl, 5 mM MgCl2, 0.1% Triton, 250 mM sucrose, 1 mM DTT) containing RNase inhibitor (1:100, NEB, M0314) and Protease Inhibitor Cocktail (1:00, Cell Signaling Technology, 5871), using a Dounce tissue grinder (Sigma-Aldrich, D9063). The homogenate was passed through a 40 μm cell strainer (Corning, 352340) and pelleted by centrifugation at 900 g for 5 min. Cells (nuclei with attached perinuclear cytoplasm) were resuspended in 1 mL staining buffer (PBS containing 0.8% BSA, RNase Inhibitor as above) and stored in Eppendorf test tubes at −80°C until FACS sorting.

### Immunostaining of neurons, astrocytes and microglia

Samples were incubated for 20 minutes in 20 μL of Fc block (Invitrogen,14-9161-73) in staining buffer. Primary antibodies were then added for 30 minutes as follows: labelled anti-NeuN for neurons (1:500; Alexa Fluor™488 conjugated, Merck, MAB377X), anti-GFAP for astrocytes (1:1000; Invitrogen, 14-9892-80), and anti-Iba1 for microglia (1:500; Abcam, ab178846). Cells were pelleted by centrifugation at 900 *g* for 5 minutes, resuspended in staining buffer and incubated for 30 minutes with secondary antibodies for neurons (1:500; Alexa Fluor™ 647 conjugated, Jackson ImmunoResearch, 711-605-152) and astrocytes (1:500; Cy3-conjugated; Jackson ImmunoResearch, 115-165-072). Cells were pelleted again as above and resuspended in staining buffer containing DAPI (1μg/mL; Santa Cruz, sc-3598). All steps were performed on ice or at 4°C.

### Immunostaining of oligodendrocytes

For oligodendrocyte isolation, nuclei were fixed and permeabilized using a specialized kit (Invitrogen, 00-5523-00), according to the manufacturer’s protocol. Anti-OLIG2 (1:100; Invitrogen, P21954) was then added in staining buffer (PBS, 0.8% BSA, RNase Inhibitor) for 1 hour. Following centrifugation and resuspension in staining buffer, cells were incubated with a secondary antibody (1:200; Alexa Fluor™ 647 conjugated, Jackson ImmunoResearch, 711-605-152) for 30 minutes. Cells were pelleted again as above and resuspended in staining buffer containing DAPI (1μg/mL; Santa Cruz, sc-3598). All steps were performed on ice or at 4°C.

### FACS-sorting

Within the DAPI-positive population, manual gates for NeuN-, IBA1-, and GFAP-positive nuclei were defined using thresholds determined from non-stained controls of the same sample (Supplementary Fig. 1e-h, j; Methods). To minimize overlap across populations, each sorting gate was refined in two stages by assessing its intersection with two other cell type populations. For example, NeuN (FITC) and IBA1 (Alexa 647) gates were first defined based on their overlap in the FITC vs the Alexa 647 space (Supplementary Fig. 1b) and then refined by excluding GFAP (Cy3)-positive events, yielding NeuN-clean and IBA1-clean populations (Supplementary Fig. 1c, d). Similarly, the initial GFAP (Cy3) gate was defined based on the overlap with the NeuN (FITC) signal (Supplementary Fig. 1c) and then refined by comparison to the IBA1 (Alexa 647)-associated signal (Supplementary Fig. 1d).

NeuN-, GFAP-, Iba1-, and OLIG2-positive cells were sorted through an 85 μm nozzle with an approximate flow rate of 8,000 events/s. Sorted populations (> 1,000 cells) were collected into tubes containing 500 μL staining buffer. Samples were collected with BD FACSAria III (BD Biosciences) and analyzed using the FCS Express 7 Software (*De Novo* Software).

### RNA extraction

Cells were centrifuged at 900 *g* for 5 minutes, resuspended in 100 μL PKD buffer (RNeasy FFPE Kit, Qiagen, 73504) and RNA was extracted (RNeasy FFPE Kit) according to manufacturer’s instructions. Concentration and RIN values were determined (Bioanalyzer 6000, Agilent Technologies), with concentrations between 0.1-12.1 ng/µL and RIN between 1.2 and 7.4. The RIN range most likely reflects varying amounts of rRNA present in the samples, in addition to possible cleavage due to the formaldehyde fixation, and not low RNA quality due to degradation.

### RT-qPCR

Synthesis of cDNA from lncRNA and mRNA was done using qScript™ cDNA Synthesis Kit (Quantabio, 95047) and qPCR was done using PerfeCTa® SYBR® Green FastMix® (Quantabio, 95072) with human-specific forward and reverse primers (Supplementary Table 15). The CFX384 Touch Real-Time PCR System (Bio-Rad) was used for quantification and the CFX Maestro software (Bio-Rad v4.1.2433.1219) for extracting Cq values. The reference gene was GAPDH. All primers were human specific.

Synthesis of cDNA from small RNAs was done using the RNA Poly(A) Tailing Kit (MCLAB, RPTK-200) and qScript Flex cDNA Synthesis Kit (Quantabio, 95049) with a proprietary oligo (dT) adaptor primer, followed by qPCR using PerfeCTa® SYBR® Green FastMix® Low ROX (Quantabio, 95074) with specific forward primers and a proprietary universal reverse primer. The CFX384 Touch Real-Time PCR System (Bio-Rad) was used for quantification and the CFX Maestro software (Bio-Rad v4.1.2433.1219) for extracting Cq values. The reference gene was hsa-let-7a-5p. All primers were human specific.

### RNA sequencing and alignment

Libraries for short RNA sequencing were constructed in two batches from 500 pg (Batch 1) and 1000 pg (Batch 2) total RNA, using the D-Plex Small RNA-seq Kit for Illumina (Diagenode, C05030001) and Single Indexes for Illumina - Set #B (Diagenode, C05030011). The small RNA fraction was separated (Invitrogen, G401004) and verified (TapeStation, Agilent Technologies), then sequenced (Illumina, 20046810 or 20046811) using the NextSeq 2000 System (Illumina) at the Center for Genomic Technologies, the Hebrew University of Jerusalem. Small RNA was aligned to miRBase 22.1 (Kozomara et al, 2019; Griffiths-Jones et al, 2007) using miRDeep (Friedländer et al, 2008).

### MicroRNA DE analysis

Raw read counts were preprocessed using DESeq2 (Love et al., 2014). Library size normalization factors were estimated using the estimateSizeFactors function to correct for sequencing depth across samples. Lowly expressed genes were filtered out by requiring counts per million (CPM) > 1 in at least 50% of samples. For the retained genes, normalized expression values were transformed as log₂ (CPM + 1).

To quantify the contribution of biological and technical factors to expression variability we used the variancePartition framework (Hoffman & Schadt, 2016). Variance components were estimated using the linear mixed model with the formula ∼(1|*ID*) + (1|*Sex*) + (1|*Batch*) + (1|*Cell. type*) + *Age*, allowing estimation of the proportion of variance explained by donor identity, sex, batch, cell type, and age.

DE analysis was then performed using the dream framework implemented in variancePartition, which models both fixed and random effects. The DE model was defined as ∼0 + *Cell. Type + Batch + *Sex* + Age +* (1|*ID*). Pairwise contrasts were computed to compare Neurons vs Microglia, Microglia vs Astrocytes, and Neurons vs Astrocytes. To identify oligodendrocyte-enriched fragments, a specific contrast comparing oligodendrocytes against all other cell types combined was constructed using the same model formulation: *Cell.typeOligodendrocyted* – 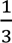 (*Cell.typeNeurons + Cell.typeMicroglia + Cell.typeAstrocytes*). Model fitting was performed with dream (Hoffman & Roussos, 2021) and limma’s eBayes functions (Ritchie et al, 2015).

### MiR mapping to cell-type-specific miR-QTLs and CREs

Annotated miR-QTLs were derived (Vattathil et al, 2025) and the cell-type-specific enhancer and promoter region annotation were determined (Nott et al, 2019). To map miR genes to cell-type-specific CREs we retrieved the genomic coordinates of all annotated miRs from miRbase 22.1 (Kozomara et al, 2019; Griffiths-Jones et al, 2007). For each miR we then extracted its chromosomal position and extended the genomic interval by 500 bases upstream and downstream, to account for minor positional uncertainty and local regulatory proximity. Each extended miR interval was then intersected with the set of open chromatin regions defined as candidate CREs (Li et al., 2023). Four major brain cell classes (neurons, astrocytes, microglia and oligodendrocytes) were defined based on the provided cell type hierarchical taxonomy (Li et al, 2023) and all corresponding fine-grained cell types were grouped under these classes.

### Single-cell RNA-sequencing analysis

The raw count matrix was downloaded from https://cellxgene.cziscience.com/collections/84ce6837-548d-4a1f-919f-0bc0d9a3952f. The raw data was normalized with scanpy’s sc.pp.normalize_total() and log-transformed with sc.pp.log1p(). The differential expression analysis was performed with sc.tl.rank_genes_groups(groupby=“class”, method=“wilcoxon”).

### IsomiR abundance analysis

Canonical and isomiR reads were aligned to miRNA precursor sequences using the miRDeep2 pipeline (Friedländer et al., 2012) by parsing the per-sample alignment outputs (.mrd) to extract isomiR-level read counts. For each precursor block, the canonical transcript was defined as the unique sequence with the highest total read count summed across all samples. Each isomiR was then classified relative to the prominent transcript of its mature unit, based on the start and end positions of its alignment, into five categories: prominent (identical start and end), 5′ overhang (start upstream of the prominent transcript, i.e., extended at the 5′ end), 5′ truncation (start downstream of the prominent transcript, i.e., recessed at the 5′ end), 3′ overhang (end downstream of the prominent transcript), and 3′ truncation (end upstream of the prominent transcript). Reads carrying coordinated shifts at both ends were assigned to several categories concurrently (e.g., 5′ truncation and 3′ overhang). Per-sample isomiR read counts were converted to counts per million (CPM) by library-size normalization, and filtered (median CPM >=10 across samples) prior to downstream analysis.

To identify individual miRNAs with cell-type-specific isomiR usage, the relative isomiR abundance was defined, per miRNA and per sample, as the ratio of isomiR CPM to canonical-transcript CPM for each of the four isomiR classes. For every miRNA and isomiR class a one-way ANOVA was performed across the four cell types, with cell type as the explanatory factor. P-values from the omnibus tests were corrected for multiple testing using the Benjamini–Hochberg procedure and miRNAs with FDR-adjusted p < 0.05 were further examined with Tukey HSD post-hoc tests to identify the cell-type pairs driving the difference (Tukey adjusted p < 0.05).

### Cell-type signal deconvolution

To infer the cellular enrichment of bulk-derived miR lists we implemented a reference-based deconvolution approach based on our cell-type-specific atlas. This was done by computing the overlap between the lists of input miRs and cell-type specific miR markers and calculating the fraction of cell-type-specific marker miRs detected in the input. To statistically assess whether an observed overlap exceeded random expectation we applied Fisher’s exact test. For each cell type, a 2 × 2 contingency table was constructed representing: (i) the number of overlapping miRs between the input and the cell type markers, (ii) the number of miRs present in the input but not among that cell type markers, (iii) the number of miRs specific to the cell type but absent from the input, and (iv) the remaining background miRs not belonging to either set. One-sided Fisher’s exact tests (alternative = ‘greater’) were used to identify significant cell type enrichments. Resulting *p-*values were corrected for multiple testing using the false discovery rate (FDR) method, yielding adjusted *p*-values.

### Elucidating age-dependent miRs with bioIB

The log-normalized miR count matrix was stored as an AnnData object. Hierarchical bioIB^75^ (described in detail in our recent paper, reference 39) was then applied using the bioIB.compress() function, with age bin (younger vs. older) specified as the signal of interest and parameters set to β = 1000, num_betas = 40, and bulk = True. For calculating the normalized mutual information (NMI) scores between metagene-based clustering and true labels of the age group (younger vs. older) and cell types, the data was clustered to 4 clusters based on the metagene profiles, using k-means. miR target genes were obtained from miRTarBase^82–84^. We included only targets that were experimentally validated in brain tissue using at least one of the following methods: HITS-CLIP, microarrays, luciferase reporter assay, Western blot, or qPCR. Gene Ontology (GO) enrichment analysis was performed using gseapy with the GO_Biological_Process_2025 library^85,86^. Significant GO terms (adjusted p ≤ 0.05) were further filtered to retain categories with at least three target genes. For improved interpretability, enriched GO terms were subsequently summarized using GO Slim subsets.

## Data availability

Raw and processed data generated in this study are available in GEO under accession number GSE314790. In addition, both the atlas and the miR cell-type deconvolution framework are available on our website for interactive exploration (https://brain-visualization.vercel.app/).

## Acknowledgements

We would like to whole-heartedly thank the neurosurgery patients who contributed to this study. We would also like to express our gratitude to Dr. Nadav Yayon, a former PhD student in the lab (currently at the Cambridge Stem Cell Institute) for developing NuNeX and for assisting with the experimental pipeline; to Dr. Ola Karmi from the Flow Cytometer Unit, at the Research Infrastructures Unit of the Hebrew University of Jerusalem, Edmond J. Safra Campus - Givat Ram (Tzabam), for her guidance and assistance in FACS sorting; and to all members of the Soreq lab for assistance and constructive input. We also thank Emmanuel Segal and Hen Shahak for their assistance in FACS sorting.

## Funding

This work was supported by the Israel Science Foundation (ISF 835/23 to H. S.), Keter Holdings, Ken Stein and William Kilberg foundations (to H.S.) and a joint grant from the Shaare Zedek Medical Center, Jerusalem (to I. P. and H. S.). Serafima Dubnov is supported by fellowships from the Azrieli, Kaete Klausner, Teva Foundations and by the Pamela and Paul Austin Research Center on Ageing.

## Competing interests

The authors report no competing interests.

## Declaration of generative AI and AI-assisted technologies in the manuscript preparation process

During the preparation of this work, the authors used Claude for polishing the text and Claude Code to assist with the code for the data analysis and figuregeneration. The authors reviewed and edited the output as needed and take full responsibility for the content of the published article.

## Notes

https://brain-visualization.vercel.app/

https://www.ncbi.nlm.nih.gov/geo/query/acc.cgi?acc=GSE314790

### Competing Interest Statement

The authors have declared no competing interest.

### Summary of Updates

*Changed the title and the abstract

