## Supplementary Figures for "Human brain microRNAs exhibit cell-type specificity and age dependence"

### Supplementary Figure 1

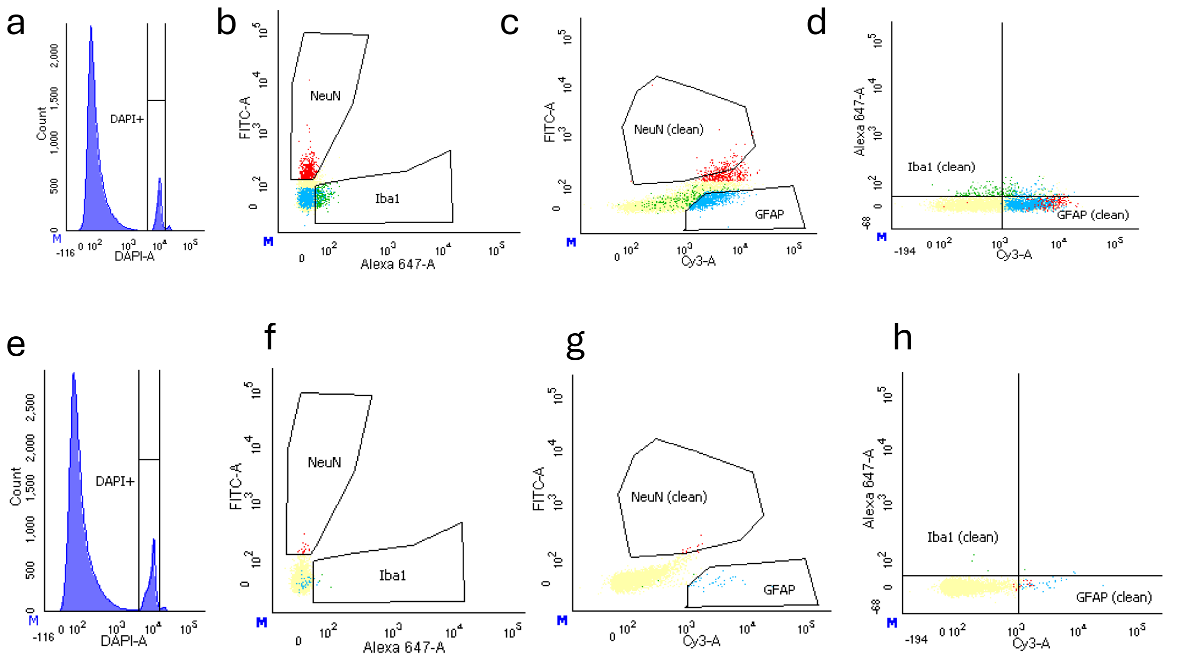


i Tube: TEST

| Population | #Events | %Parent | %Total |
|---|---|---|---|
| All Events | 67,273 | #### | 100.0 |
| Scatter gate | 61,967 | 92.1 | 92.1 |
| SSC | 61,327 | 99.0 | 91.2 |
| FSC | 61,308 | 100.0 | 91.1 |
| DAPI+ | 5,691 | 9.3 | 8.5 |
| NeuN | 438 | 7.7 | 0.7 |
| Q1-2 | 2 | 0.5 | 0.0 |
| Q2-2 | 436 | 99.5 | 0.6 |
| Q3-2 | 0 | 0.0 | 0.0 |
| Q4-2 | 0 | 0.0 | 0.0 |
| NeuN (clean) | 115 | 26.3 | 0.2 |
| Iba1 | 249 | 4.4 | 0.4 |
| Iba1 (clean) | 95 | 38.2 | 0.1 |
| Q2-1 | 138 | 55.4 | 0.2 |
| Q3-1 | 7 | 2.8 | 0.0 |
| Q4-1 | 9 | 3.6 | 0.0 |
| GFAP | 960 | 16.9 | 1.4 |
| Q1-3 | 0 | 0.0 | 0.0 |
| Q2-3 | 80 | 8.3 | 0.1 |
| Q3-3 | 4 | 0.4 | 0.0 |
| GFAP (clean) | 876 | 91.2 | 1.3 |

j Tube: NC

| Population | #Events | %Parent | %Total |
|---|---|---|---|
| All Events | 100,000 | #### | 100.0 |
| Scatter gate | 91,466 | 91.5 | 91.5 |
| SSC | 90,209 | 98.6 | 90.2 |
| FSC | 90,196 | 100.0 | 90.2 |
| DAPI+ | 10,244 | 11.4 | 10.2 |
| NeuN | 15 | 0.1 | 0.0 |
| Q1-2 | 8 | 53.3 | 0.0 |
| Q2-2 | 7 | 46.7 | 0.0 |
| Q3-2 | 0 | 0.0 | 0.0 |
| Q4-2 | 0 | 0.0 | 0.0 |
| NeuN (clean) | 8 | 53.3 | 0.0 |
| Iba1 | 5 | 0.0 | 0.0 |
| Iba1 (clean) | 2 | 40.0 | 0.0 |
| Q2-1 | 3 | 60.0 | 0.0 |
| Q3-1 | 0 | 0.0 | 0.0 |
| Q4-1 | 0 | 0.0 | 0.0 |
| GFAP | 35 | 0.3 | 0.0 |
| Q1-3 | 0 | 0.0 | 0.0 |
| Q2-3 | 2 | 5.7 | 0.0 |
| Q3-3 | 0 | 0.0 | 0.0 |
| GFAP (clean) | 33 | 94.3 | 0.0 |

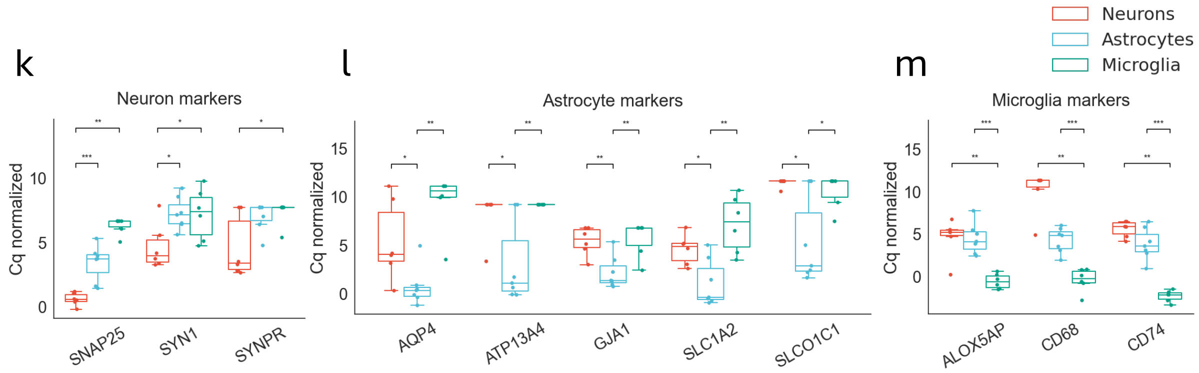

**Supplementary Figure 1**. a) Histogram featuring the DAPI signal of FACS events, showing the defined DAPI-positive gate, separating single cells (middle peak) from debris (left peak, low signal levels) and doublets (right peak, higher signal level). b) Scatter plot showing Alexa 647 (IBA1) and FITC (NeuN) signals of the FACS events from the DAPI-positive gate shown in (a), with NeuN- and IBA1-positive gates. c) Scatter plot showing Cy3 (GFAP) and FITC (NeuN) signals of the DAPI-positive FACS events, with the refined NeuN-clean and GFAP-positive primary gates. d) Scatter plot showing the Cy3 (GFAP) and Alexa 647 (IBA1) signals of DAPI-positive FACS events, with the refined Iba1-clean and GFAP-clean gates ensuring minimal mutual overlap. e) Histogram of DAPI fluorescence for FACS events in a representative negative Control (NC) sample stained with DAPI only. f) Scatter plot showing Alexa 647 (IBA1) vs. FITC (NeuN) signals for FACS events in the NC sample stained with DAPI only. g) Scatter plot showing Cy3 (GFAP) vs. FITC (NeuN) signals for DAPI-positive FACS events in the NC sample stained with DAPI only. h) Scatter plot of Cy3 (GFAP) versus Alexa 647 (IBA1) signals for DAPI-positive gated events in the NC sample, showing refined IBA1-clean and GFAP-clean gates with minimal mutual overlap. i) FACS table quantifying the proportions of events assigned to each gate in the Test sample (e-h). j) Same as (i) for the NC sample (a-d). k) Box plots of GAPDH-normalized qPCR Cq values for the neuron markers SNAP25, SYN1 and SYNPR across all three cell types. l) Box plot as in g for the astrocyte markers AQP4, ATP13A4, GJA1, SLC1A2 and SLCO1C1. m) Box plot as in g for the microglial markers AQP4, ATP13A4, GJA1, SLC1A2 and SLCO1C1. Outliers were removed for each gene's cell type box using the standard boxplot definition, excluding values lying below Q1 – 1.5×IQR or above Q3 + 1.5×IQR. $^*p \leq 0.05$, $^{**}p \leq 0.01$, $^{***}p \leq 0.001$ (non-parametric Mann–Whitney U test).

### Supplementary Figure 2

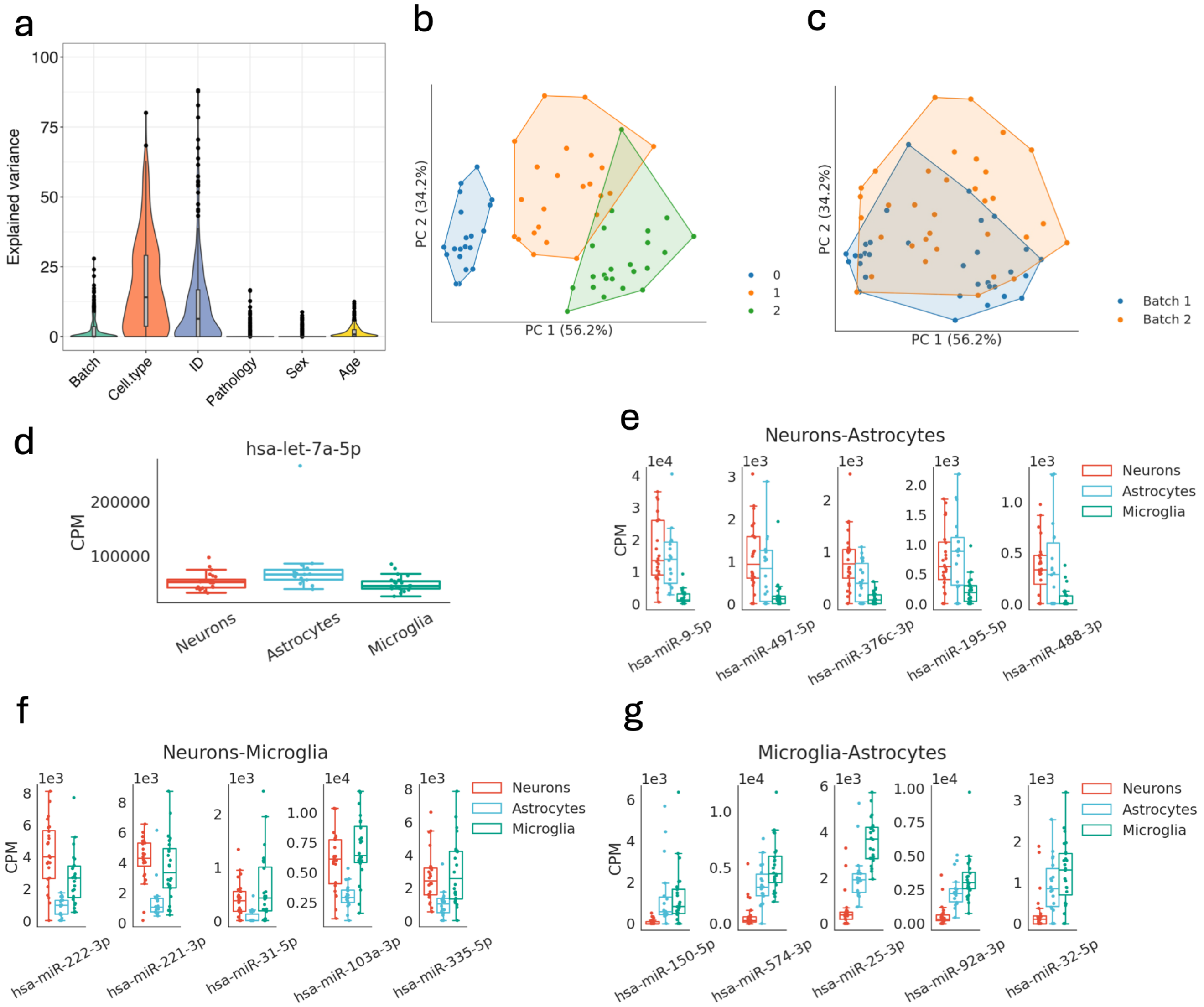


**Supplementary Figure 2.** a) Violin plot generated with the *variancePartition* package, showing the proportion of explained variance in miR expression profiles attributed to Batch, Cell type, Patient ID, Pathology (meningioma/metastasis), Sex, and Age. b) PCA of miR expression profiles colored by k-means cluster assignments. Convex hulls outline each cell-type cluster. c) PCA of miR expression profiles colored by Batch. Convex hulls outline each cell-type cluster. d) Box plot showing the expression of hsa-let-7a-5p across the three cell types. Due to its uniform levels, it was selected as a house-keeping gene in miR qPCR measurements. e) Box plots of the top five miRs (ranked by mean log fold change) significantly elevated in both neurons and astrocytes relative to microglia. f) Box plots of the top five miRs significantly elevated in both neurons and microglia relative to astrocytes. g) Boxplots of the top five miRs significantly elevated in both microglia and astrocytes relative to neurons.

### Supplementary Figure 3

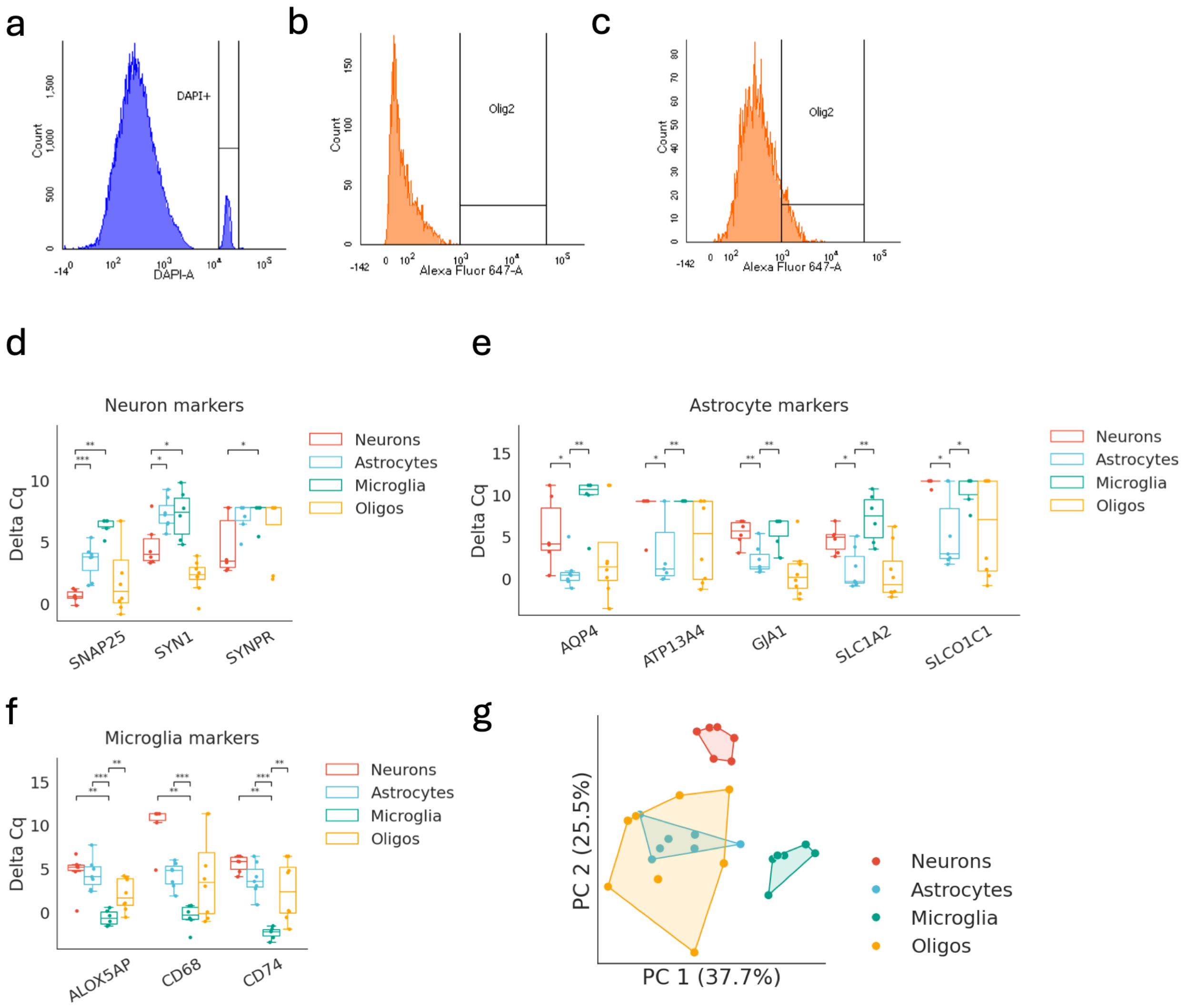


**Supplementary Figure 3.** a) Histogram of DAPI fluorescence for FACS events in a representative sample stained with DAPI and OLIG2 for oligodendrocyte sorting. b) Histogram of Alexa 647 (OLIG2) fluorescence in a representative NC sample stained with DAPI only. c) Same as (b) for the same sample  stained with both DAPI and OLIG2. d) Box plots showing GAPDH-normalized qPCR Cq levels of neuronal markers *SNAP25*, SYN1 and *SYNPR* across sorted populations (neurons, astrocytes, microglia, and oligodendrocytes). e) Same as (d) for astrocyte markers *AQP4, ATP13A4, GJA1, SLC1A2 and SLCO1C1*. f) Same as (d) for microglia markers *ALOX5AP, CD68* and *CD74*. *p ≤ 0.05, **p ≤ 0.01, ***p ≤ 0.001 (non-parametric Mann–Whitney U test). g) PCA of GAPDH-normalized Cq values across all samples, colored by cell type. Convex hulls outline each cell-type cluster.

### Supplementary Figure 4

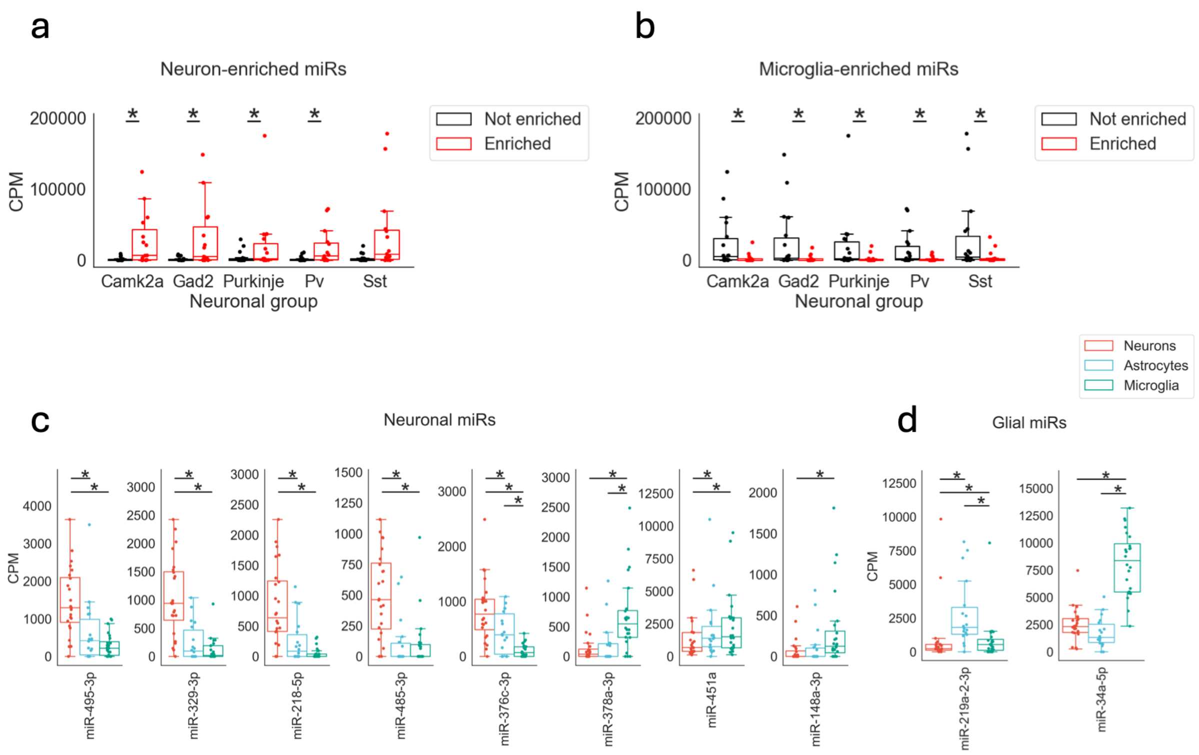


**Supplementary Figure 4.** a) Box plots of CPM-normalized miR levels across the neuronal subtypes profiled by He et al.[1], with miRs stratified into those enriched in neurons in our atlas (red) versus those not enriched in neurons (black). b) As in (a), but with miRs stratified into those enriched in microglia in our atlas (red) versus those not enriched in microglia (black). (c, d) Box plots of CPM-normalized miR levels across three cell types in our atlas (neurons, astrocytes, microglia), with miRs split into two subplots based on Merienne et al.[2]: (c) miRs identified as neuron-enriched relative to glia; (d) miRs identified as glia-enriched relative to neurons. *p ≤ 0.05, **p ≤ 0.01, ***p ≤ 0.001 (Mann–Whitney U test).

### Supplementary Figure 5

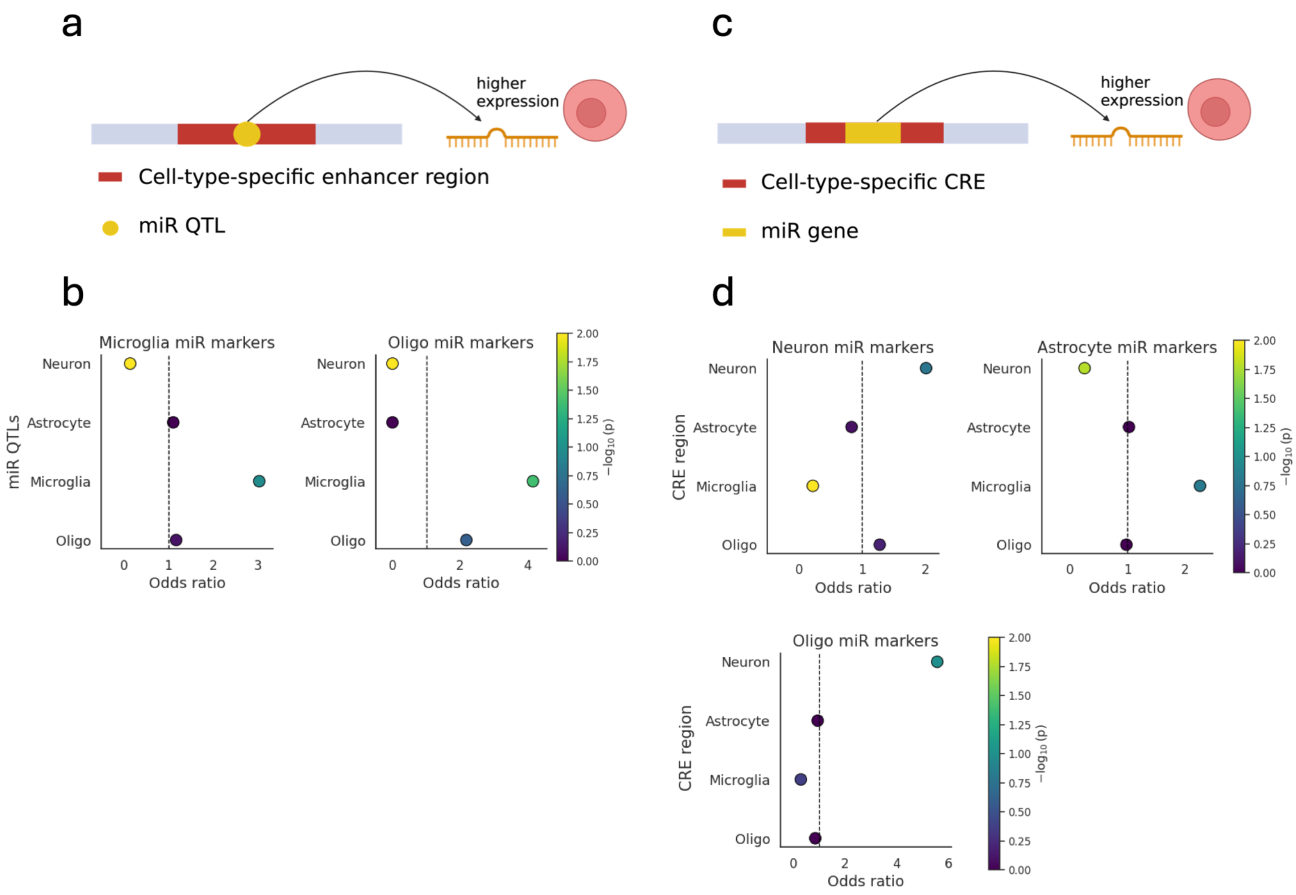


**Supplementary Figure 5.** a) Schematic representation of a miR-QTL residing in a cell-type-specific enhancer region, contributing to the cell-type-specific expression of the associated miR. Created with BioRender. b) Dot plots showing Fisher's exact test results (alternative = two-sided) for the association between miRs enriched in microglia (left) and oligodendrocytes (right), and the associated miR-QTLs residing in cell-type-specific enhancer regions. Y-axis depicts cell type, X-axis positions reflect the odds ratio of each association, and dot colors depict $-\log_{10}$ p-values (unadjusted). c) Schematic representation of a miR gene located within a cell-type-specific cis regulatory element (CRE) (red) that contributes to cell-type-specific expression of the miR.. Created with BioRender. d) Dot plots showing Fisher's exact test results (alternative = two-sided) for associations between transcriptomically enriched miRs in neurons (top left), astrocytes (top right), and oligodendrocytes (bottom left), and miRs residing within cell-type-specific cis-regulatory elements (CRE). Y-axis depicts cell type, X-axis positions reflect the odds ratio of each association, and dot colors depict $-\log_{10}$ p-values (unadjusted).

### Supplementary Figure 6

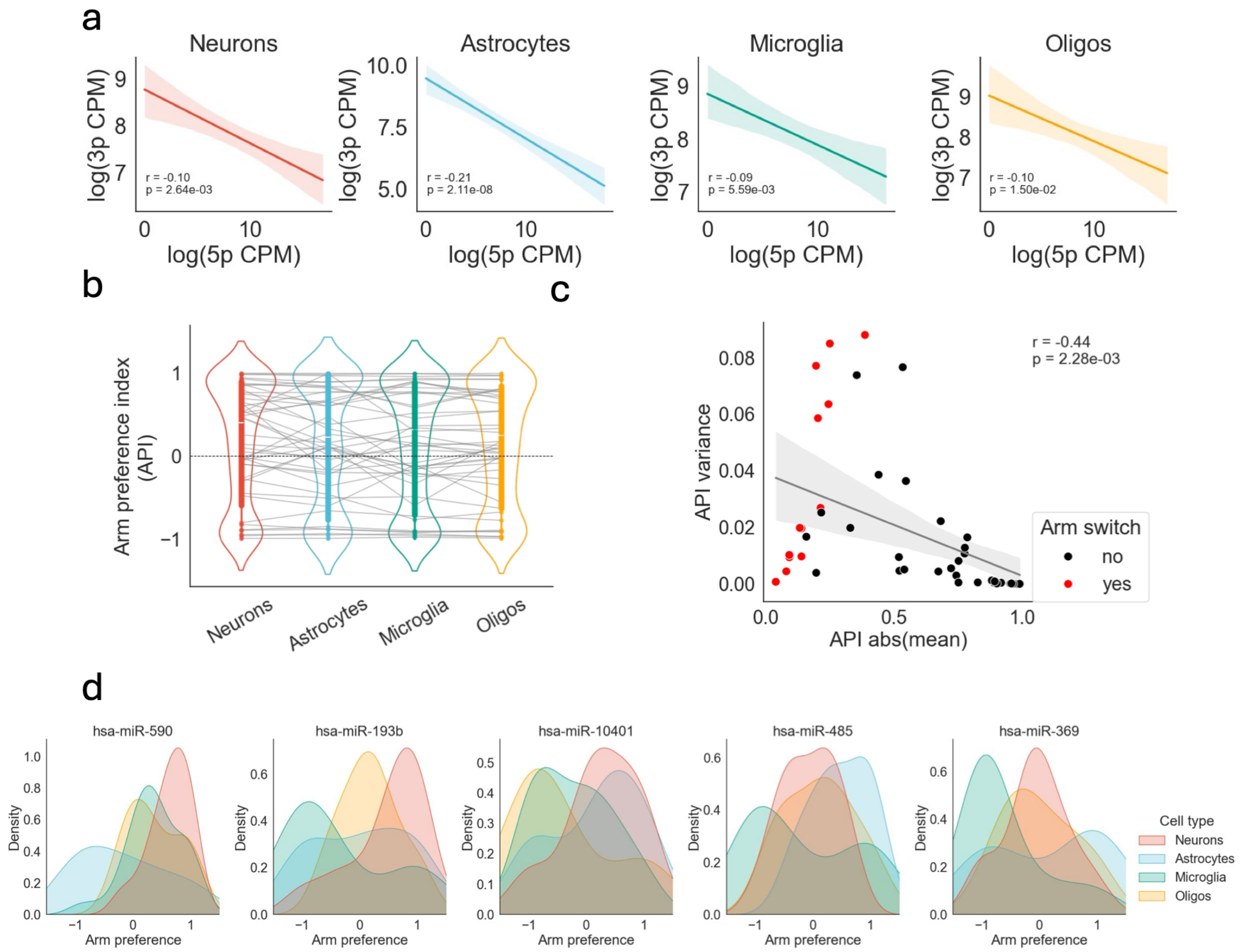


**Supplementary Figure 6.** a) Regression plots of log(CPM+1) expression of 5′-arm versus 3′-arm miRNAs derived from the same precursor, shown separately for each cell type. Pearson's R and the corresponding p-value are indicated on each panel. b) Violin plots of the Arm Preference Index (API) distribution in each cell type; grey lines connect the same miRNA precursor across cell types. c) Regression plot showing a significant negative correlation between the absolute mean API and the API variance across cell types; miRNA precursors that switch their dominant arm in a cell-type-dependent manner are highlighted in red. d) Kernel density estimates (KDE) of the API distribution across cell types for five miRNA precursors with cell-type-dependent arm dominance (API variance > 0.05).

### Supplementary Figure 7

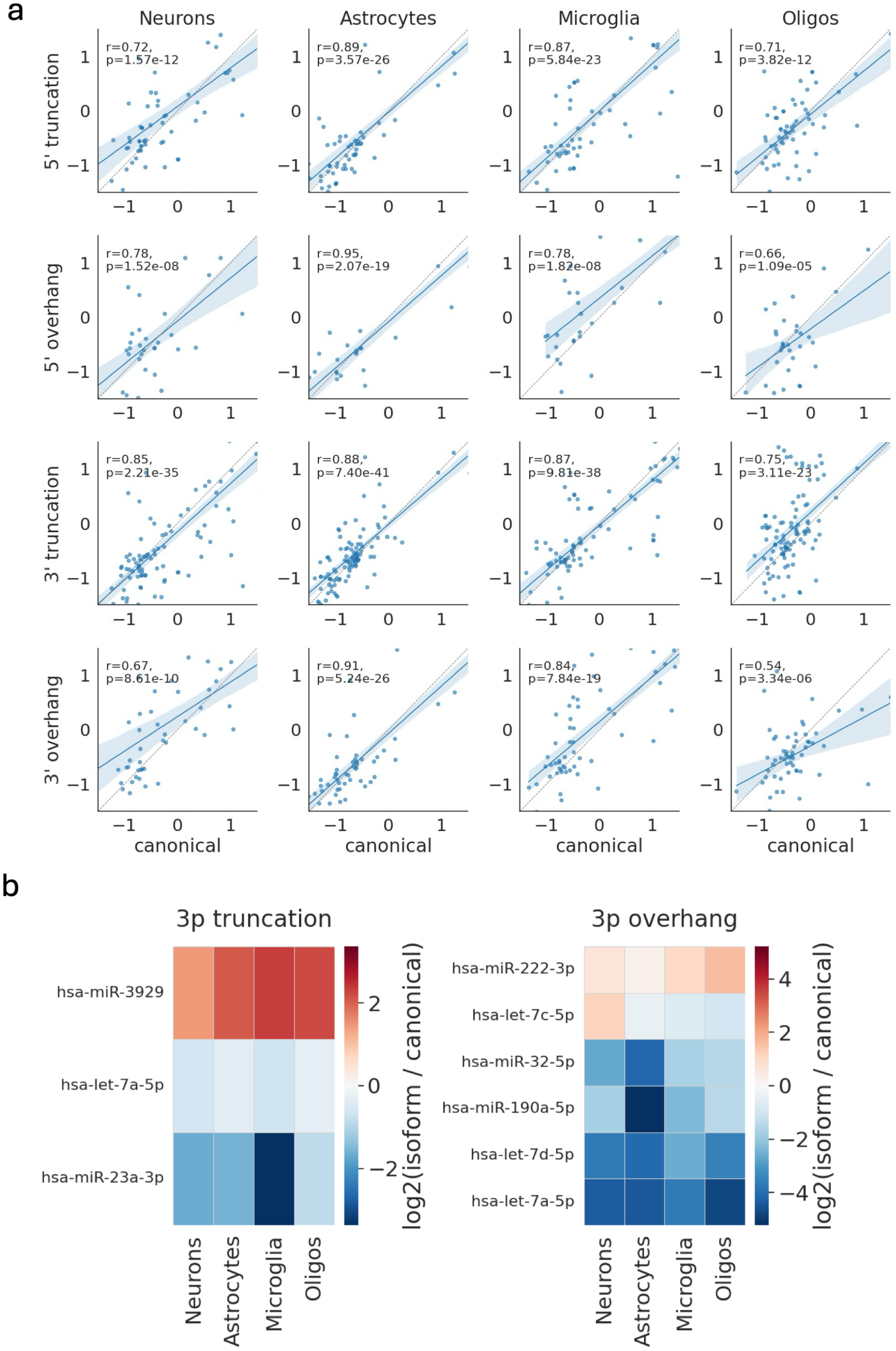

**Supplementary Figure 7.** a) Regression plots of isomiR mean normalized levels (z-scores) versus those of the corresponding canonical miRNAs, with isomiR classes as rows (5′ truncation, 5′ overhang, 3′ truncation, 3′ overhang) and cell types as columns. Pearson's R and the corresponding p-value are indicated on each panel. b) Heatmap of the log-transformed CPM difference between each isomiR and its canonical miRNA, restricted to isomiRs whose abundance differs significantly across cell types (Methods).

#### Supplementary Figure 8

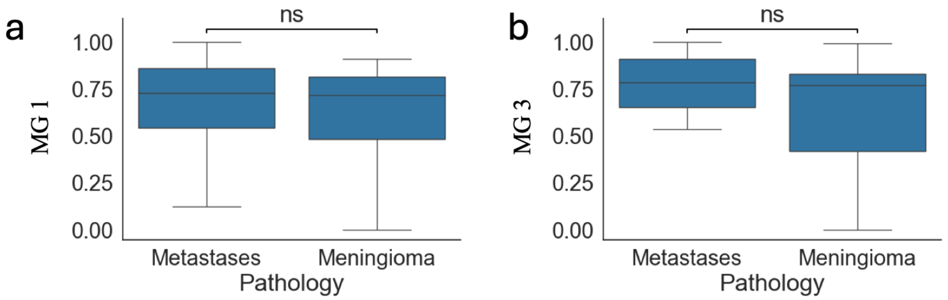


**Supplementary Figure 8.** Box plots of the normalized expression levels of metagene 1 (a) and metagene 3 (b) in samples from patients with brain metastases versus meningioma. ns, not significant ($p > 0.05$, Mann–Whitney U test).
